# Chunking as a rational strategy for lossy data compression in visual working memory

**DOI:** 10.1101/098939

**Authors:** Matthew R. Nassar, Julie C. Helmers, Michael J. Frank

**Author notes:** Corresponding Author: Matthew R. Nassar, Department of Cognitive, Linguistic and Psychological Sciences, Brown University, Providence, RI 02912-1821.

## Abstract

The nature of capacity limits for visual working memory has been the subject of an intense debate that has relied on models that assume items are encoded independently. Here we propose that instead, similar features are jointly encoded through a “chunking” process to optimize performance on visual working memory tasks. We show that such chunking can: 1) facilitate performance improvements for abstract capacity-limited systems, 2) be optimized through reinforcement, 3) be implemented by center-surround dynamics, and 4) increase effective storage capacity at the expense of recall precision. Human performance on a variant of a canonical working memory task demonstrated performance advantages, precision detriments, inter-item dependencies, and trial-to-trial behavioral adjustments diagnostic of performance optimization through center-surround chunking. Models incorporating center-surround chunking provided a better quantitative description of human performance in our study as well as in a meta-analytic dataset, and apparent differences in working memory capacity across individuals were attributable to individual differences in the implementation of chunking. Our results reveal a normative rationale for center-surround connectivity in working memory circuitry, call for re-evaluation of memory performance differences that have previously been attributed to differences in capacity, and support a more nuanced view of visual working memory capacity limitations: strategic tradeoff between storage capacity and memory precision through chunking contribute to flexible capacity limitations that include both discrete and continuous aspects.

## Introduction

People are limited in their capacity to retain visual information in short-term memory; however, the exact nature of this limitation is hotly debated (Luck & Vogel, 2013; Ma, Husain, & Bays, 2014; Wei, Wang, & Wang, 2012). Competing theories have stipulated that capacity is constrained by either a discrete item limit (e.g., a fixed number of “slots”) or by the distribution of a flexible “resource” across relevant visual information (Bays & Husain, 2008; Wei et al., 2012; Zhang & Luck, 2008). In their simplest form, these competing theories are both philosophically distinct and statistically identifiable, but experimental evidence has been mixed, with some studies favoring each theory and the best-fitting computational models incorporating elements of each (Almeida, Barbosa, & Compte, 2015; Bays & Husain, 2008; Bays, Catalao, & Husain, 2009; Cowan & Rouder, 2009; Chris Donkin, Tran, & Nosofsky, 2013a; Christopher Donkin, Nosofsky, Gold, & Shiffrin, 2013b; Rouder et al., 2008; van den Berg, Awh, & Ma, 2014; van den Berg, Shin, Chou, George, & Ma, 2012; Zhang & Luck, 2008; 2009; 2011). Experimental support for both theories has emerged from delayed report working memory tasks, in which subjects are asked to make a delayed report about a feature (e.g. color) of a single item that was briefly presented as part of a multi-item stimulus display (Bays & Husain, 2008; Wilken & Ma, 2004; Zhang & Luck, 2011). In particular, as the number of items to be retained increases, visual working memory reports tend to become less precise, as predicted by resource models, and more likely to reflect guessing, as predicted by slots models (Fougnie, Suchow, & Alvarez, 2012; Luck & Vogel, 2013; Ma et al., 2014; van den Berg et al., 2012; 2014).

While the competing classes of visual working memory models have evolved substantially over the past decade, the mathematical formalizations of each have relied on assumptions about what is, and should be, stored in working memory. Thus, an open question with potentially broad implications is what *should* and *do* people store in memory during performance of the standard delayed recall tasks, and how do deviations from the standard assumptions affect our understanding of memory capacity? To this end, recent work has highlighted the ability of people to optimize memory encoding and decoding processes by pooling information across memoranda to enhance performance under different regimes (Brady & Alvarez, 2011; 2015; Brady, Konkle, & Alvarez, 2009; Lew & Vul, 2015; Orhan & Jacobs, 2013; Sims, Jacobs, & Knill, 2012; Wei et al., 2012). Specifically, people can integrate prior information to improve memory report precision (Bays et al., 2009; Brady & Alvarez, 2011) and, when stimuli are redundant, lossless compression strategies can be used to efficiently encode them (Bays et al., 2009; Brady et al., 2009; Zhang & Luck, 2008). These strategies can improve memory performance, but only to the extent to which features of upcoming memoranda are predicted by previously observed stimulus statistics. Since memoranda in standard working memory tasks are unpredictable and randomly distributed by design, such strategies cannot improve and may actually impede performance in standard tasks (Bays et al., 2009; Orhan & Jacobs, 2014; Zhang & Luck, 2008). However, while memoranda in these tasks are not compressible in the “lossless” sense, it is still possible that people might employ more fast and frugal techniques to reduce memory storage requirements at a small but acceptable cost to task performance.

Here we explore this possibility and show that people should, could, and do implement a lossy form of data compression that sacrifices information about subtle differences in the feature values of memoranda in order to improve overall task performance. We do so using an inter-related set of computational models across different levels of analysis, such that we can constrain our understanding of the compression algorithm using both computational notions of how information should be compressed and mechanistic notions of how biological circuits could implement this compression. We probe our own and published empirical data to test key predictions of these models. This study thus involves four related components:

1. *Normative and behavioral analysis.* We begin with an information-theoretic analysis of how features of memoranda *should* be stored to maximize task performance in an abstract memory-limited system. We show that that under high memory load conditions, it is advantageous to jointly encode (chunk) a blended representation of similar features and only separately encode (partition) features if they are sufficiently dissimilar. This strategy can be effectively implemented by setting a criterion for partitioning features based on dissimilarity, where the appropriate criterion can be learned based on trial feedback (binary reward) and increases with memory load. We show that human subject behavior in a delayed report working memory task conforms to predictions from this form of adaptive chunking and optimization thereof via reward feedback.
2. *Mechanistic implementations of chunking in a biophysical network model.* Given that behavioral data accorded with predictions from the normative model, we next examined how such selective chunking *could* be implemented in a biophysical memory system. Established recurrent neural network models have shown that perceptually similar memoranda can be merged together via attractor “bump collisions” (Wei et al., 2012). However, these simulations showed that such an architecture leads to indiscriminate chunking and – due to lateral inhibition – increased forgetting of non-chunked items, leading to poor performance (Wei et al., 2012). We show that this issue can be remedied by adopting a more biologically motivated center-surround inhibition connectivity that effectively partitions dissimilar color representations; however, it does so at the cost of inter-item repulsions, which reduce the effective precision of memory reports (Almeida et al., 2015).
3. *Algorithmic model of center-surround dynamics.* To systematically explore the impact of center-surround mediated chunking at the behavioral level, we created a parsimonious (i.e., minimal parameter) model that incorporates the key features necessary for effective chunking afforded by the biophysical model without explicitly modeling the temporal dynamics or biophysical properties. We show that center-surround dynamics facilitate improved memory recall at the expense of precision, and capture previously unexplained qualitative patterns of bias and precision in human memory reports across stimulus arrays.
4. *Quantitative model fitting.* Finally, we fit behavioral data from individual subjects to show that chunking and center-surround dynamics improve fit relative to state-of-the-art models, and can account for considerable performance differences across subjects, with better-performing subjects best fit by models with more inclusive chunking policies. We validate these findings in a meta-analytic dataset to show that chunking improves quantitative model fits across tasks, offers an alternative explanation to changes in precision with set size, and accounts for individual differences in working memory task performance under conditions of high memory load.

## Results

Visual working memory capacity is typically measured using either delayed report or change detection tasks (Bays & Husain, 2008; Wilken & Ma, 2004; Zhang & Luck, 2011). Here we focus on the former, as they have provided nuanced information about the shape of memory distributions and have formed the basis for competing models of capacity limitation (Fougnie et al., 2012; Luck & Vogel, 2013; Ma et al., 2014; van den Berg et al., 2012).

Specifically, we consider a delayed report color reproduction task that requires storage of color and orientation information (figure 1). Each trial consists of three core stages: stimulus presentation, delay, and probe. During stimulus presentation, five oriented colored bars are displayed simultaneously. During the subsequent delay, the screen is blanked, requiring short-term storage of color and orientation information. During the probe stage, a single oriented bar is displayed (in gray) and the participant is required to reproduce the color that had been associated with that orientation in the preceding stimulus array.

**Figure 1:**
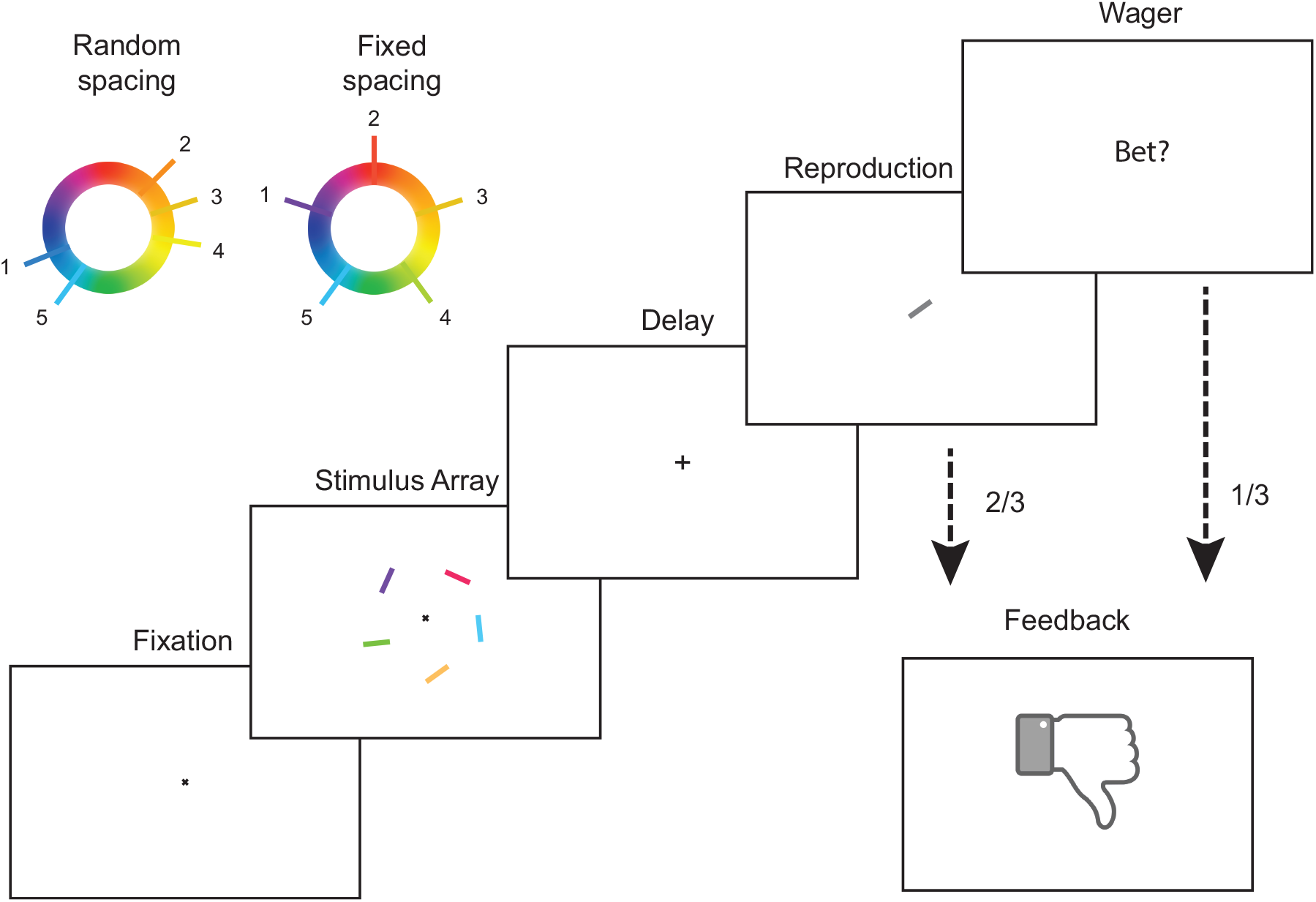
Delayed report color reproduction task. Each trial begins with central fixation for 500 ms, followed by stimulus presentation for 200 ms. Stimuli consist of five colored and oriented bars evenly distributed around a circle subtending 4 degrees of visual angle and centered on the point of fixation. Stimulus presentation is followed by a 900 ms delay, after which a single oriented bar is displayed centrally. The subject is required to report the color associated with the bar with the probed orientation in the previous stimulus array. After confirming the report, the subject receives feedback dependent on whether the absolute magnitude of the reproduction error was greater or less than a fixed threshold. Stimulus colors on any given trial are selected either: 1) randomly and independently as is standard in such tasks (random spacing; upper left) or 2) ensuring uniform spacing on the color wheel so as to minimize within-array color similarity (fixed spacing).

### Information theoretic analysis of what should be stored in working memory

To first understand whether, in principle, information encoding could be optimized in this task, we developed a limited-capacity system for memory storage in which colors and orientations are represented with binary words (figure 2). We build on the now classical work of George Miller by conceptualizing working memory capacity limitation in terms of a strict limit on information storage, in bits (G. A. Miller, 1956). The precision with which a color is stored depends on the number of binary digits (bits) used to represent that color: a single bit can be used to specify a half of the color wheel, a second bit can be added to specify a quarter of the color wheel, and so on (figure 2A,). Capacity limitations within such a system can be easily implemented as a fixed limit on the number of bits stored during the delay period. These bits can be used to represent the individual colors in the target array, by, for example, dividing them evenly among the available targets (figure 2B).

Alternatively, multiple similar colors could be jointly represented with a single binary word that is then linked to multiple orientations (figure 2C). An intuitive advantage of the second encoding strategy is that reducing the number of binary color words increases the number of bits available to represent each word, potentially offsetting the biased encoding of the chunked items by better representing each encoded color.

**Figure 2:**
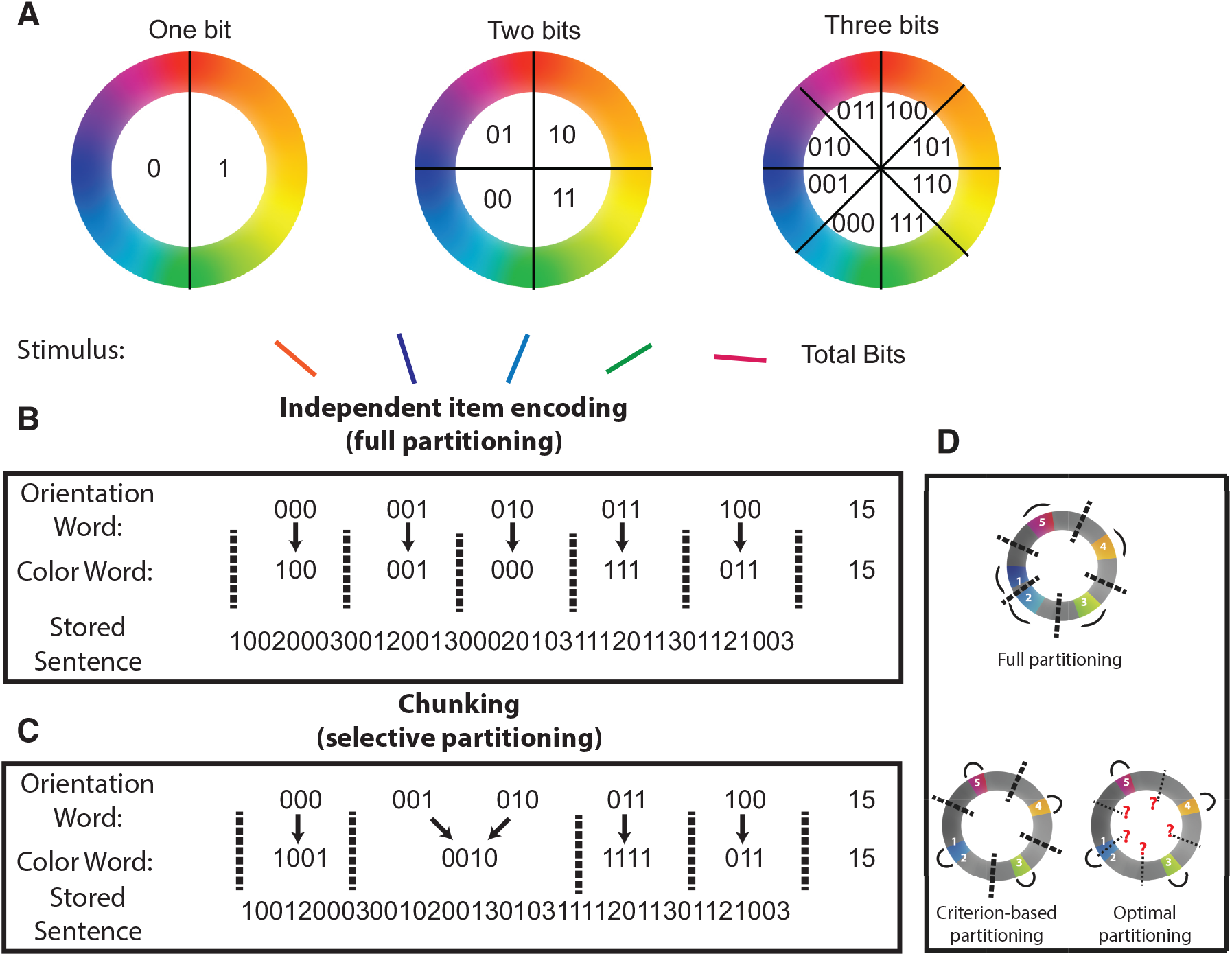
Binary encoding model of visual working memory. In order to formalize capacity limitations, it is useful to consider an abstract model of working memory that stores features in binary words. **A**: Each color can be described by a binary word of fixed length, where the number of digits in the word determines the storage precision. **B & C**: Stimulus arrays can be stored by linking ordered pairs of color and orientation words. Capacity limitations are modeled by a fixed limit on the length of the resulting “sentence” comprised of color and orientation words separated by word termination symbols (2/3 for color/orientation words, respectively). **B:** One strategy for storing ordered pairs involves alternating sequences of color and orientation words, such that each color is “partitioned” from all other colors (dotted lines separating color representations) and linked to a single orientation. **C**: Another strategy for storage would be to link two or more orientations to a single color by removing a partition (chunking). This reduces the number of colors that need to be stored, and thus increases the number of bits allotted to each color. **D**: Full partitioning (top) involves placing a partition between each set of colors such that each color is represented independently. Criterion-based partitioning sets a partition between each set of colors that are separated by a greater distance than the partitioning criterion. Optimal partitioning examines all partitioning patterns for a given stimulus array and selects the partitioning pattern that would achieve the lowest theoretical error magnitude**. Colors/Arcs in each model reflect stored representations of a particular stimulus array (actual stimuli labeled with numbers) and thick/thin lines indicate actual/potential partitions. Note that in this case, actual partitions selected by optimal partitioning do not differ from those selected by the criterion-based partitioning model.**

To test this potential advantage quantitatively, we examined task performance of three models that differ in how they “chunk” feature information. The standard model employs independent item encoding, through which each color is partitioned, or represented separately, from all other stored colors (full partitioning; figure 2B&D). We also consider a fully optimal model that considers all possible partitioning patterns and stores information using the combination of partitioning and chunking (figure 2C&D) that would lead to the best theoretical performance for each specific stimulus array, given the possibility that any item could be probed (optimal partitioning). Although determining the optimal partitioning pattern for each array is computationally expensive, it is well approximated by a simple and efficient heuristic, using a single criterion for partitioning, according to the separation of two targets in color space (criterion-based partitioning; fig 2D). For this model, chunking was parameterized by a “partitioning criterion” that defines the minimum distance between two colors required for independent representation. If the distance between two colors is smaller than the partitioning criterion, the colors are represented as a single “chunk”. Thus, a partitioning criterion of zero indicates that all items are represented independently, whereas a partitioning criterion of π indicates that all item colors will be chunked together (i.e. represented by a single binary word). Performance of all models was assessed for delayed recall tasks ranging from easy (2 items) to difficult (8 items) and using both continuous and discrete assumptions regarding the distribution of item information.

Performance of the criterion-based model depended on partitioning criterion as a function of task difficulty (figure 3). For easier tasks with few items to encode, the model’s memory buffer was large enough to store each item independently with a reasonable number of bits, such that increasing the partitioning criterion beyond zero was detrimental to task performance (two targets; figure 3a [dark line]). However, for harder tasks, in which storing each item with high precision was not possible due to limited buffer size, performance was best for moderate partitioning criterions that allow for joint representation of similar, but not dissimilar, colors (eight targets; figure 3a [light line]). To better understand how chunking interacts with set size to affect performance, we compared the performance of the best criterion-based partitioning model to that of a full partitioning model across different task difficulties. Across task difficulties, there was a monotonic relationship between the number of targets and the performance advantage of both fully optimal and criterion-based chunking models over full partitioning (figure 3B; compare orange and green lines). Furthermore, the performance of the best criterion-based partitioning was nearly identical to that of the optimal partitioning model (figure 3B; compare green and yellow lines). Notably, the number of colors stored by criterion-based and optimal partitioning models saturated around four with increasing set size, highlighting that the improved performance of these models comes from chunking similar colors into a single representation that is linked to multiple orientations. Moreover, this result suggests that even though it is possible in these models to store more items independently with less precision, it is more advantageous to restrict the number of stored representations during standard delayed recall tasks (figure 3C).

Set-size dependent performance advantages of chunking were also relatively insensitive to modeling assumptions regarding the nature of storage constraints. While the results described above were generated under the assumption of a divisible resource framework (figure 3A-C), comparable results were attained when model behavior was simulated using binary words that roughly correspond to the slots + averaging framework (figure 3D-F). Thus, the performance advantages offered by criterion-based chunking are robust to the nature of the actual capacity limitation.

### Adaptation of partitioning criterion via reinforcement learning

The advantages of chunking discussed above were presented for the best partitioning criterion, which differed as a function of task demands, begging the question of how a human subject would know to use this criterion. We thus examined whether the partitioning criterion could be optimized on a trial-to-trial basis via reinforcement learning to improve performance by allowing the partitioning criterion to be adjusted on each trial according to the chunking (total number of chunks) and reward feedback (thresholded binary feedback) from the previous trial (see Methods). The resulting model increased the partitioning criterion, and thus chunking, after a trial in which it chunked colors and achieved better-than-average performance. This led partitioning criterions to increase rapidly towards a load-dependent asymptote (figure 3G). Trial-to-trial increases in the partitioning criterion corresponded to rapid improvements in overall task performance, as measured by average absolute error (figure 3H). These improvements in task performance were concomitant with reductions in the total number of color chunks that the model attempted to store (figure 3I). Thus, chunking can be optimized across conditions through reinforcement learning to improve performance and reduce effective storage requirements in higher memory load contexts.

Given the performance advantages offered by criterion-based chunking and the efficiency with which it could be learned and implemented, we next sought to identify diagnostic predictions made by the model. One key difference between the criterion-based and full partitioning models is that the performance of the former depended heavily on the specific distribution of the colors within each stimulus array. If the colors were randomly and independently sampled, which is the standard method in such tasks, chunking offered large advantages. In contrast, the advantages of chunking were considerably smaller when colors were uniformly distributed in color space to maximize color separation (fixed spacing; figure 1 inset; compare the solid and dashed lines in figure 3E-F). It is noteworthy that the prediction of performance decrements under fixed spacing conditions is completely opposite to that made by a prominent active maintenance model of working memory, a point that we address in detail with a biologically motivated model of chunking in a later section (Wei et al., 2012).

**Figure 3:**
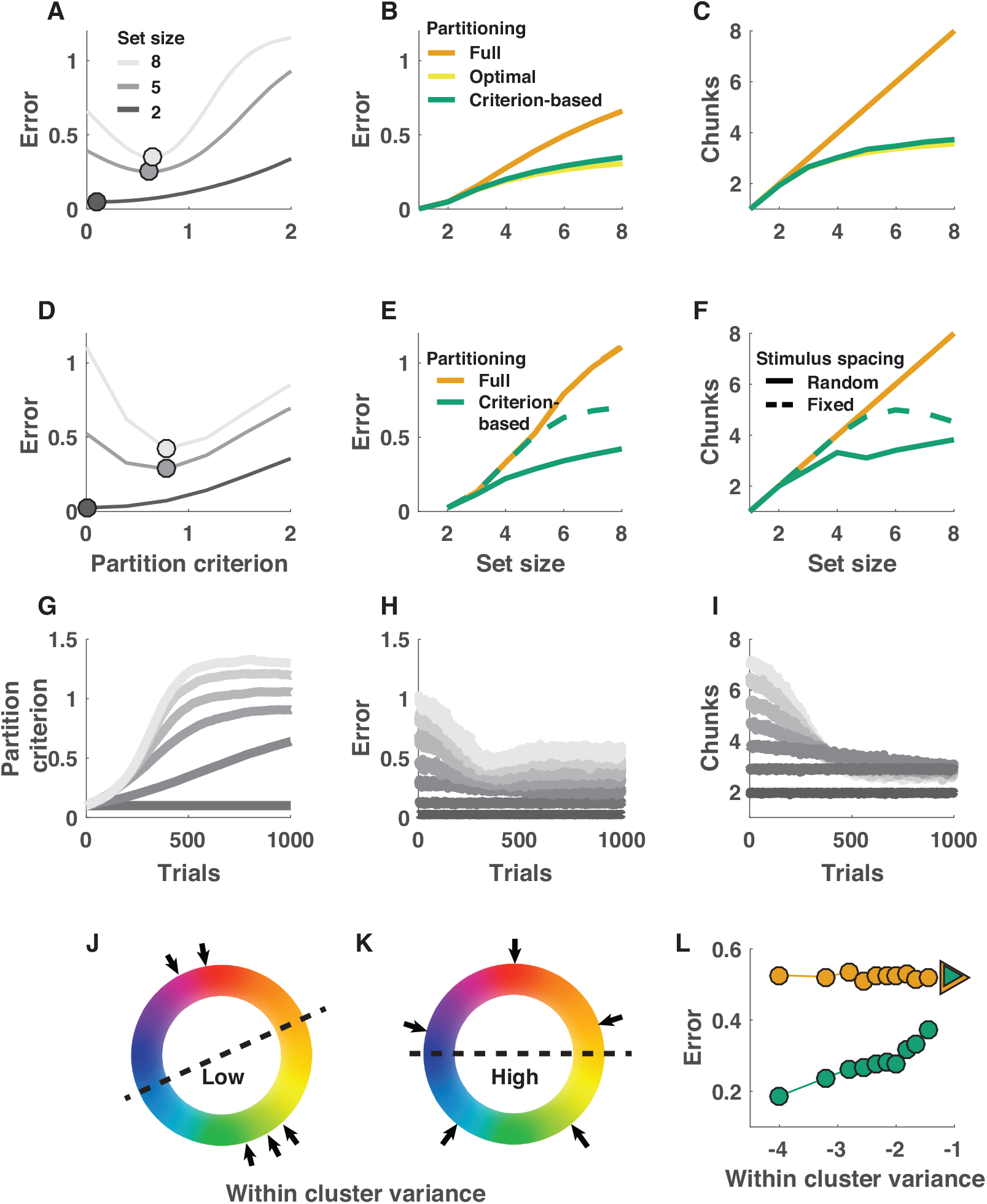
Chunking improves memory performance and can be achieved through trial-to-trial adjustments of partitioning criterion. **A-C**: Criterion-based chunking confers memory performance advantages and reduces feature storage requirements under resource assumptions. **A:** Mean absolute error (ordinate) for theoretical performance of a binary encoding model on delayed report tasks of varying set size (grayscale) across all possible partitioning criterions (abscissa; 0 = all colors stored independently). **B:** Model error (ordinate) increases as a function of set size (abscissa) for three partitioning strategies: 1) fully partitioned (model always stores all targets independently), 2) optimal partitioning (model considers all possible partitions for each stimulus array and uses the best), 3) criterion-based partitioning (chunking and partitioning is determined by best criterion from **A**). Error increases more shallowly for optimal and criterion-based partitioning strategies that employ strategic chunking. **C:** Total number of chunks requiring storage (ordinate) increases as a function of set size (abscissa) for all three models, but saturates near 4 items for optimal and criterion-based chunking models. **D-F:** Performance advantages of criterion-based chunking hold for binary word storage, analogous to “slots + averaging.” **D-F** are analogous to **A-C** except that panels E and F show model performance separately for randomly spaced and fixed spaced stimulus arrays (solid and dotted lines, respectively) and do not include an “optimal partitioning” model, as computing it would be computationally inefficient under this framework. **G-I:** Appropriate partitioning criterions can be learned through reinforcement. **G**: Adjusting the partitioning criterion through reinforcement learning (see Methods) leads simulated criterions (ordinate) to increase over trials (abscissa) in a manner that scales with set size (grayscale; 2 = darkest, 8 = lightest). Adjustments in criterion lead to reduced errors **(H)** and decrease the “chunks” that require storage **(I)**. **J-L:** Chunking selectively benefits performance on trials in which colors are most tightly clustered. Within-cluster variance provides a measure of feature clustering within a stimulus array, with low values indicating more clustering (**J**) and high values indicating less clustering (**K**). Performance of the best chunking model, but not the non-chunking model, depends on the clustering of individual stimulus arrays, as assessed through within-cluster variance. Mean absolute error is plotted for stimulus arrays grouped in bins of within-cluster variance for criterion-based chunking (green) and fully partitioned (orange) models. Triangles reflect the same values computed for fixed spacing trials, in which stimulus features were minimally clustered (as depicted in **K**).

To better characterize the statistical properties of the specific arrays that enabled better performance in the chunking models, we computed the within-cluster variance (WCV) as a metric of how efficiently each stimulus array could be chunked (see Methods). Target arrays with tightly clustered colors have low WCV, whereas target arrays with distantly spaced colors have high WCV (figure 3J&K). The performance of chunking models depended monotonically on WCV, with the smallest errors achieved on low WCV target arrays (figure 3L), supporting the notion that chunking advantages are achieved by more efficient representation of similar colors through joint encoding.

Taken together, the results from the binary encoding model suggest that 1) selectively chunking similar feature values can improve performance in working memory tasks, 2) performance improvements from selective chunking increase with target number and are mitigated by uniformly spacing feature values, 3) performance of chunking models depends monotonically on WCV, and 4) chunking behavior can be learned through reinforcement of successful chunking behaviors. In summary, the binary encoding model clarifies why, and under what conditions, chunking could benefit working memory performance in standard tasks. In the next section, we test whether performance of human subjects in a delayed recall task conforms to the predictions of the criterion-based chunking model.

### People are more accurate and confident when remembering clustered stimulus arrays

To directly test key predictions made by the binary encoding model, we collected behavioral data from human participants in the task described in figure 2. The critical manipulation in the task is that the colors in some trials were uniformly spaced on the color wheel (fixed spacing) whereas the colors in interleaved trials were randomly and independently sampled from the color wheel (random spacing). This manipulation allowed us to test the prediction of the binary coding model that more clustered stimulus arrays lead to better performance (figures 3E&I). To examine the potential for adaptive learning of chunking, we provided reward feedback to subjects on each trial (by comparing error magnitude to a fixed threshold, either π/3 or π/8 in separate groups). The task also required subjects to wager about the accuracy of their choices (post-decision) on one third of trials. These task features allowed us to test the prediction that chunking behaviors are adjusted from trial-by-trial through reinforcement learning (see figure 3G&H) and to determine whether participants were aware of any performance bonuses attributable to chunking.

In accordance with behavioral optimization through selective chunking, participants were more accurate and confident when presented with randomly spaced stimuli. Subject accuracy, as assessed using the same error threshold procedure used to determine feedback, was greater on random spacing trials than on fixed spacing trials (figure S1A; t = 4.4, p < 10e-4). This accuracy improvement was more prevalent for subjects that experienced a liberal accuracy criterion (low precision, absolute error < π/3) than for those that experienced a conservative accuracy criterion (high precision, absolute error < π/8; two sample t-test for group difference: t = -2.5, p = 0.02, Bayes Factor = 4.8) suggesting the improvement may have been more pronounced for moderate errors that were interpreted differently across the groups depending on the accuracy criterion (error < π/3 & error > π/8). Participants also wagered more frequently on random spacing than fixed spacing trials, suggesting that they were cognizant of the conferred performance advantage (figure S1B; t = 3.1, p = 0.003). Subjects tended to gauge their wagering behavior reasonably well, with subjects who achieved higher levels of accuracy also betting more frequently (figure S1C; Pearson’s rho =0.68, p <10e-6). Furthermore, individual differences in the adjustment of wagering as a function of color spacing configuration correlated with the change in accuracy that subjects experienced across the configurations (figure S1D; Pearson’s rho =044, p = 0.002). Taken together, these data suggest that subjects experienced and were aware of performance advantages for randomly spaced stimuli, but that the extent of these advantages differed across individuals.

To better understand these performance advantages, we tested the extent to which trial-to-trial accuracy and confidence scores depended on stimulus clustering within the randomly spaced stimulus arrays. Specifically, we computed within-cluster variance (WCV) for each color array (as for the binary model) to evaluate whether this clustering statistic could be used to predict subjects’ accuracy of, and confidence in, color reports. As predicted by chunking models (figure 3I), subjects were more accurate for low WCV trials; performance on high WCV trials was similar to that in the fixed spacing configuration (figure 4A). Furthermore, subject wagering also decreased monotonically with WCV, such that betting behavior on the highest WCV (least clustered) color arrays was similar to that on fixed spacing trials (figure 4B).

**Figure 4:**
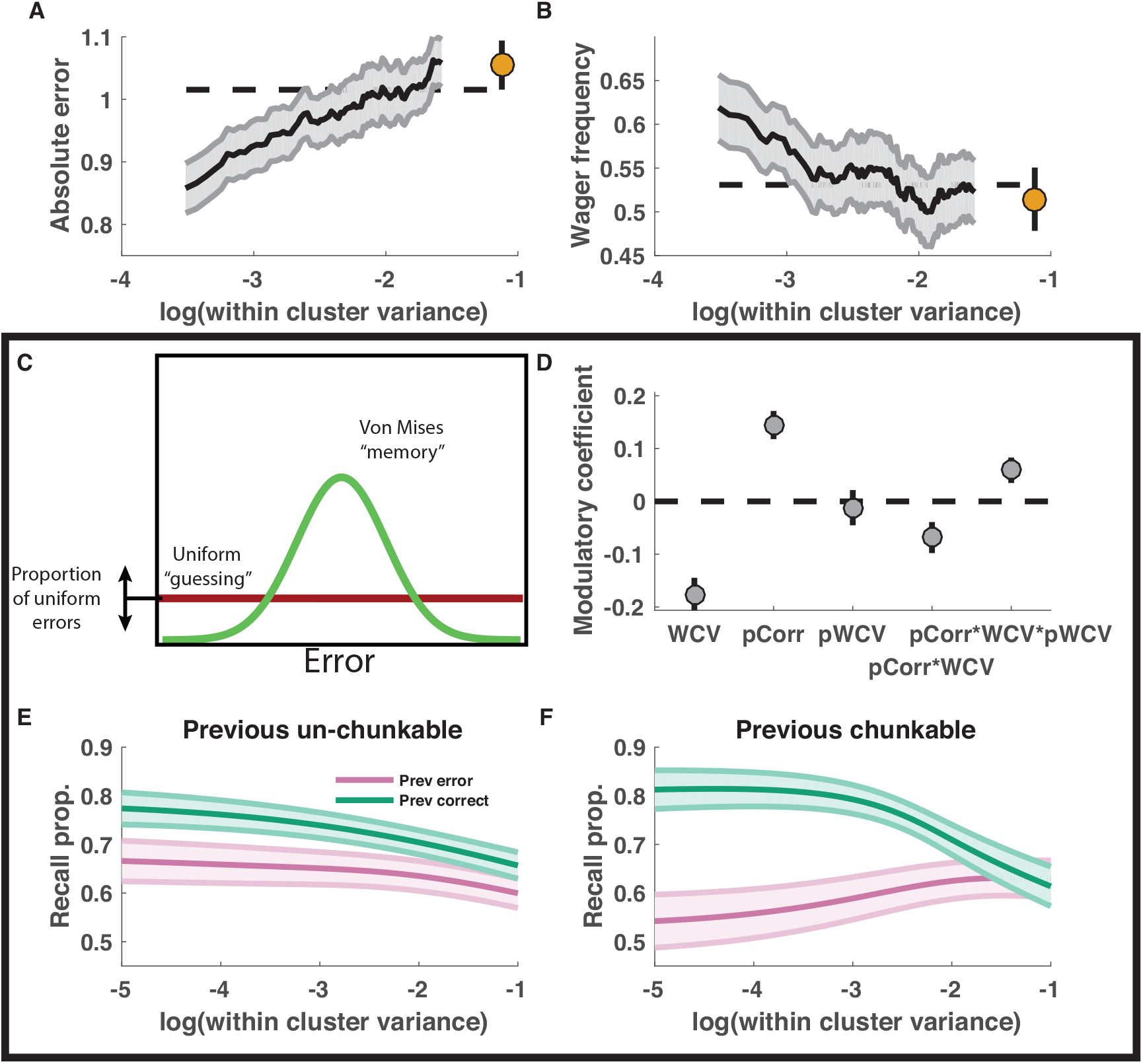
Memory recall and confidence are enhanced for clustered stimulus arrays and adjusted according to trial feedback in accordance with model predictions. **A&B:** Memory performance and confidence increase with stimulus clustering. Mean absolute error magnitude (**A**) and high wager frequency (**B**) were computed per subject in sliding bins of within-cluster variance (larger values = decreased stimulus clustering) for random (lines) and fixed spacing conditions (points). Lines and shading reflect mean and SEM across subjects. **C-F**: Mixture model fits reveal recall benefit of stimulus clustering and hallmark of feedback-driven criterion adjustments. **C**: Subject data were fit with a mixture model that considered reports to come from a mixture of processes including 1) a uniform “guess” distribution, 2) a “memory+binding” distribution centered on the color of the probed target, and 3) a “binding error” distribution including peaks at each non-probed target [not shown]. Additional terms were included in the model to allow the recall probability to vary as a logistic function of stimulus clustering, recent feedback, and their interaction. **D-F:** Recall probability was modulated by feedback and stimulus clustering in a manner suggestive of trial-to-trial adjustments of chunking. Mean/SEM coefficients across subjects for each modulator of recall (log within-cluster variance (WCV), previous trial feedback (pCorr), previous trial log within-cluster variance (pWCV), pCorr*WCV and pCorr*WCV*pWCV) are represented from left to right as points/lines. Multiplicative interaction terms were included to capture the form of criterion adjustments that were used to facilitate criterion learning in the binary encoding model (Fig 3G-I). **E&F:** Recall probability of best-fitting descriptive models plotted as a function of the log within-cluster variance for the current trial and divided according to previous feedback (color) and the log within-cluster variance from the previous trial [E: pWCV =-1, F: pWCV=-5]. Lines/shading reflect mean/SEM across subjects. Feedback effects are consistent with reinforcement-learning as implemented in the binary encoding model: when chunking clustered stimulus arrays is rewarded with positive feedback, it is reinforced, leading to selective performance improvements for clustered stimulus arrays on the subsequent trial.

### Performance advantages are not due to binding errors

An alternative explanation of these effects is that subjects exhibited higher performance in low WCV trials simply because they committed binding errors (Bays et al., 2009), mistaking one color for a nearby related color. To address this issue, we applied a generalized linear model (GLM) to the binary accuracy and confidence data. In particular, we included the distances between the target color and each non-probed color as nuisance variables to determine whether the apparent WCV effects could be better explained by a tendency to report non-probed colors, which are often closer to the target color for more clustered stimulus arrays. When this model was applied separately to subject accuracy and accuracy of reports simulated from a mixture model that included binding errors, coefficients for WCV were negative in fits to subject data but negligible when fit to simulated data, suggesting that performance improvements mediated by WCV were not simply a reflection of binding errors (figure S2; subject accuracy β = 0.028, t = 5.3, p < 10e-6). Furthermore, when the same model was fit to post-decision wagers, coefficients for WCV took similarly negative values, suggesting that subjects were aware of the performance advantages that they gained from the clustered stimulus arrays (β = -0.049, t=-5.7, p<10e-7).

### Trial-by-trial adjustment in accordance with reinforcement learning

The GLM also included terms to account for other factors that could potentially affect task performance, including feedback from previous trials. Positive feedback on the previous trial was predictive of higher accuracy and confidence for the current trial, in a manner consistent with trial-by-trial behavioral adjustment (figure S2, “correct (t-1)” coefficient; accuracy β = 0.017, t = 5.0, p < 10e-6; confidence β = 0.026, t = 4.9, p < 10e-5). This predictive relationship was unlikely to be driven by autocorrelation in performance, as such an explanation should also predict that confidence measurements relate to accuracy on future trials, and this relationship was not evident in the data (figure S2, “correct (t+1)” coefficient; confidence β = 0.0017, t = 0.3, p = 0.75). Despite seemingly robust feedback-driven effects, overall performance improvements across the session were somewhat modest, as evidenced by a relatively small positive coefficient for a term in the GLM relating block number to accuracy (figure S2, “block” coefficient; accuracy β = 0.007, t = 2.3, p = 0.02, Bayes Factor = 2.3). Thus, the GLM results suggest that subjects gained a performance advantage for clustered target arrays, modulated behavior in response to previous feedback, and improved slightly over the course of task sessions. Below we provide a more specific test of whether subjects were adjusting chunking strategies in accordance with the reinforcement learning strategy employed in the binary encoding model.

To better understand how working memory fidelity depends on stimulus clustering and feedback history, we extended a basic mixture model of subject memory reports (figure 4C; (Bays et al., 2009; Zhang & Luck, 2008)). The model considers memory reports to come from a mixture distribution that includes a “memory” component centered on the probed color, a “guessing” component uniformly distributed across color space and a “binding error” component that assumes reports are centered on non-probed target colors (not shown in figure; (Bays et al., 2009; Zhang & Luck, 2008)). We allowed the proportion of recall (1-guessing) to vary as a function of the key factors that should affect performance if subjects are optimizing chunking. Across subjects, the probability of recall increased with stimulus clustering, as assessed by within-cluster variance (figure 4D; t = -5.7, p < 10e-6). Notably, performance improvements due to stimulus clustering were observed even for stimulus arrays in which the probed target color was dissimilar to the cluster of non-probed target colors (figure S3), as predicted by the binary encoding model (i.e., where chunking increases resources and probability of encoding of other items). Together with the GLM results above, these findings rule out alternative explanations in which WCV effects would arise simply by mistaking one color for another one nearby, and instead support our thesis that clusters of stimuli can be stored jointly to conserve representational space.

Furthermore, additional coefficients provided evidence that people adjusted chunking strategies online as a function of reward feedback in a manner similar to that used to optimize performance in the binary encoding model (figure 3G). In particular, in our reinforcement learning implementation, the partitioning criterion was adapted based on the amount of chunking in the previous trial and the concomitant reward feedback, and it selectively contributed to performance improvements for the most clustered stimulus arrays (figure 3H-F). To explore the possibility that people adjust chunking in a similar way, we included additional terms in the mixture model to allow recall probability to vary as a function of previous trial feedback (pCorr), proxies for previous and current trial clustering (pWCV, WCV), and their predicted interactions (see Methods). The best-fitting coefficients revealed an overall recall bonus on trials following correct feedback (pCorrect: t = 5.4, p < 10e-5), but also that the magnitude of this bonus was greater for trials in which stimuli were more clustered (pCorrect * WCV: t = -2.1, p = 0.04, Bayes Factor = 1.6) and for trials in which the level of stimulus clustering matched that of the previous trial (pCorrect * WCV * pWCV: t = 2.1, p = 0.04, Bayes Factor = 1.6; figure 4D). Consistent with optimization of chunking via reinforcement learning, these interactions capture a tendency toward larger feedback-driven changes in task performance when both the current and previous trial color arrays were highly clustered (figure 4E&F). Taken in the context of our model, this suggests that when subjects are rewarded for successfully chunking a stimulus array, they are more apt to repeat this chunking on the subsequent trial. Moreover, these strategic adjustments rule out an obligatory alternative mechanism in which chunking occurs regardless of task demands.

In sum, our abstract model of a memory-limited system predicted performance advantages and trial-to-trial adjustments associated with selective chunking that were subsequently validated in empirical data from human subjects performing a working memory task. Nonetheless, the abstract system that we explored does not mimic the processing architecture that is used by the brain, leaving open the question of how selective chunking might be achieved using biological hardware. In the next section we consider this question in depth, by attempting to endow a biologically inspired active maintenance model of working memory with the computational elements necessary to achieve the chunking-based performance bonus predicted by the abstract model and observed in data from human subjects.

### Center-surround dynamics as a mechanism for chunking and partitioning

The brain is thought to implement visual working memory in neural networks that include individual neurons tuned to specific visual features and capable of maintaining representations in the absence of input (Curtis & D’Esposito, 2003; Fuster & Jervey, 1981; Goldman-Rakic, 1995; E. K. Miller & Cohen, 2001; Warden & Miller, 2007). In computational models, the ability of a neural network to maintain feature representations in the absence of inputs depends critically on the recurrent connections between neurons (Barak, Sussillo, Romo, Tsodyks, & Abbott, 2013; Durstewitz & Seamans, 2002; Kilpatrick, Ermentrout, & Doiron, 2013; Murray et al., 2014; X. J. Wang, 1999). In particular, persisting feature representations, like those of the colors in our task, are facilitated by local excitatory connections between similarly tuned neurons and by broad inhibition between neurons (X. J. Wang, 1999). Importantly, for simplicity, the most common variant of such models includes uniformly weighted connections for inhibitory neurons (figure 5A). While this model exhibits merging of color representations and hence provides a promising mechanism for chunking, it produces promiscuous merging of individual color representations (bumps), large and unwieldy bumps of population activity, and due to uniform lateral inhibition, forgetting of further colors (figure 5C; (Wei et al., 2012)). Thus in a sense, such bump collisions are analogous to the chunking implemented in our more abstract binary encoding model, yet they lack the selectivity necessary to mediate performance optimization, and indeed, predict the opposite pattern of performance than seen empirically, with worse performance for randomly spaced arrays and improved performance for fixed arrays (Wei et al., 2012)

**Figure 5:**
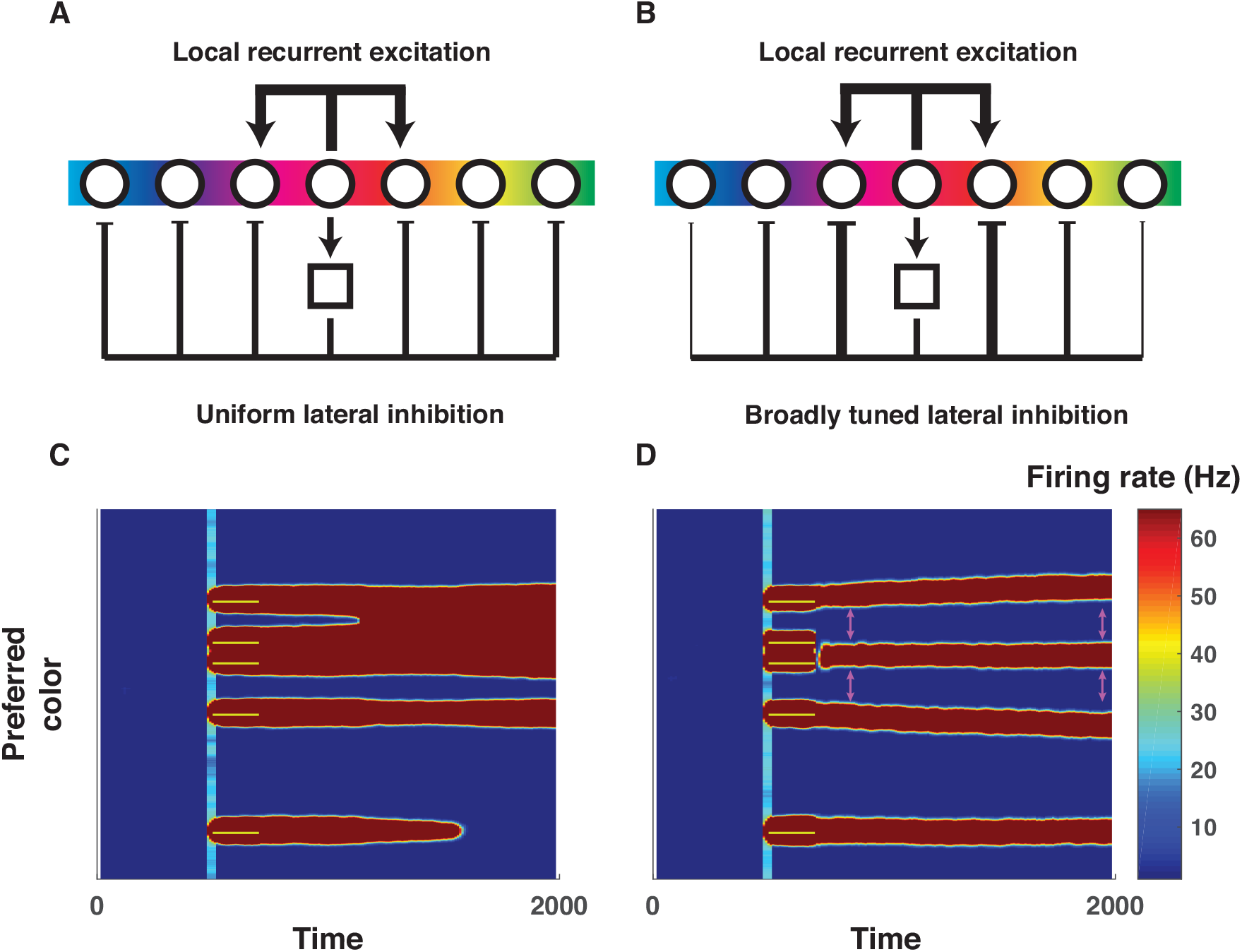
Center-surround connectivity as a mechanism to support chunking and partitioning operations needed to optimize working memory storage. **A&B)** Local recurrent excitation and lateral inhibition are critical for active working memory maintenance in biologically plausible neural networks (Almeida et al., 2015; Wei et al., 2012). However, the exact form of lateral inhibition has been varied across studies, with the most common version employing uniform inhibition across the entire population of tuned excitatory neurons (**A**, (Wei et al., 2012)) whereas others employ broadly tuned inhibition such that similarly tuned excitatory neurons indirectly exert stronger inhibitory forces on one another (**B**, (Almeida et al., 2015)). **C)** Simulated firing rates (redder colors indicate greater firing) of a population of color tuned neurons using the connectivity architecture described in panel **A** performing a working memory task (ordinate reflects neural tuning preference; abscissa reflects time in milliseconds; yellow bars indicate 200 ms color inputs delivered in a fixed pattern across network architectures). As described by Wei and colleagues, bumps of neural activity sometimes collide, producing “merged” representations (e.g., top activity bump in panel C), a possible mechanism for chunking. However, also as described by Wei and colleagues, collisions are somewhat indiscriminate and can increase overall population firing, which in turn can lead to collapse of other activity bumps (e.g., bottom activity bump) and hence forgetting. **D)** Simulated firing rates from the same population of neurons for the same task, but using center-surround connectivity (i.e., broadly tuned inhibition). Note that the closest bumps of activity are selectively chunked (e.g., second and third bump from top), but the tuned inhibition effectively partitions more distantly separated representations (e.g., the top from the second and third) and prevents forgetting of unrelated items. A related consequence of the tuned inhibition is that partitioned representations exert repulsive forces on one another during the delay period (see differences in separation of activity bumps at pink arrows). Thus, tuned inhibition affords selective partitioning of representations, but changes representations through inter-item repulsion.

We considered whether other patterns of connectivity would remedy this issue. Notably, physiological data suggest that neural responses within such networks obey center-surround receptive field architectures that are present throughout the visual system (Hubel & Wiesel, 1959; 1965), are supported by lateral connectivity (Ben-Yishai, Bar-Or, & Sompolinsky, 1995; Kohonen, 1982; Somers, Nelson, & Sur, 1995), and predict biases previously observed in visual working memory reports (Almeida et al., 2015; Kiyonaga & Egner, 2016). We thus altered the Wei et al. model to include broadly tuned inhibition in accordance with center-surround recurrence, whereby Feedback inhibition is stronger for neurons with similar color tuning (figure 5B). In this case, recurrent excitation promotes merging of nearby color representations, but tuned inhibition prevents the merged representation from expanding in width, thereby preventing it from 1) suppressing other activity bumps through an overpowering lateral inhibition leading to forgetting, and 2) indiscriminately merging with other activity bumps (figure 5D). Indiscriminate merging is prevented through repulsive forces exerted by each activity bump on neighboring representations, which enable separation of dissimilar representations (see widening gap noted by pink arrows). Thus, center-surround recurrence provides a dynamic analog to the selective chunking necessary for optimization in the binary encoding model; nearby representations that share excitatory recurrent connections are merged into a single representation (chunking) whereas more distantly separated representations are repulsed by tuned inhibition to effectively partition moderately dissimilar color representations (partitioning).

This center-surround connectivity profile promotes complex item interactions that can be summarized by a “difference of Gaussians” function that mediates the attraction and joint representation of similar colors and the repulsion of dissimilar ones (figure 6A; yellow shading). If the two stored colors are similar enough to promote mutual recurrent excitation, each represented color will experience biased excitation in the direction of its neighboring color, and eventually the two color “bumps” will merge to form a single representation (figure 6A) (Wei et al., 2012). In contrast, if the stored colors are separated beyond the narrowly tuned recurrent excitation function, mutual recurrent inhibition will dominate, leading to a net repulsion of color representations from one another (Felsen, Touryan, & Dan, 2005; Kiyonaga & Egner, 2016), which can serve as a selective partition (figure 6A).

**Figure 6:**
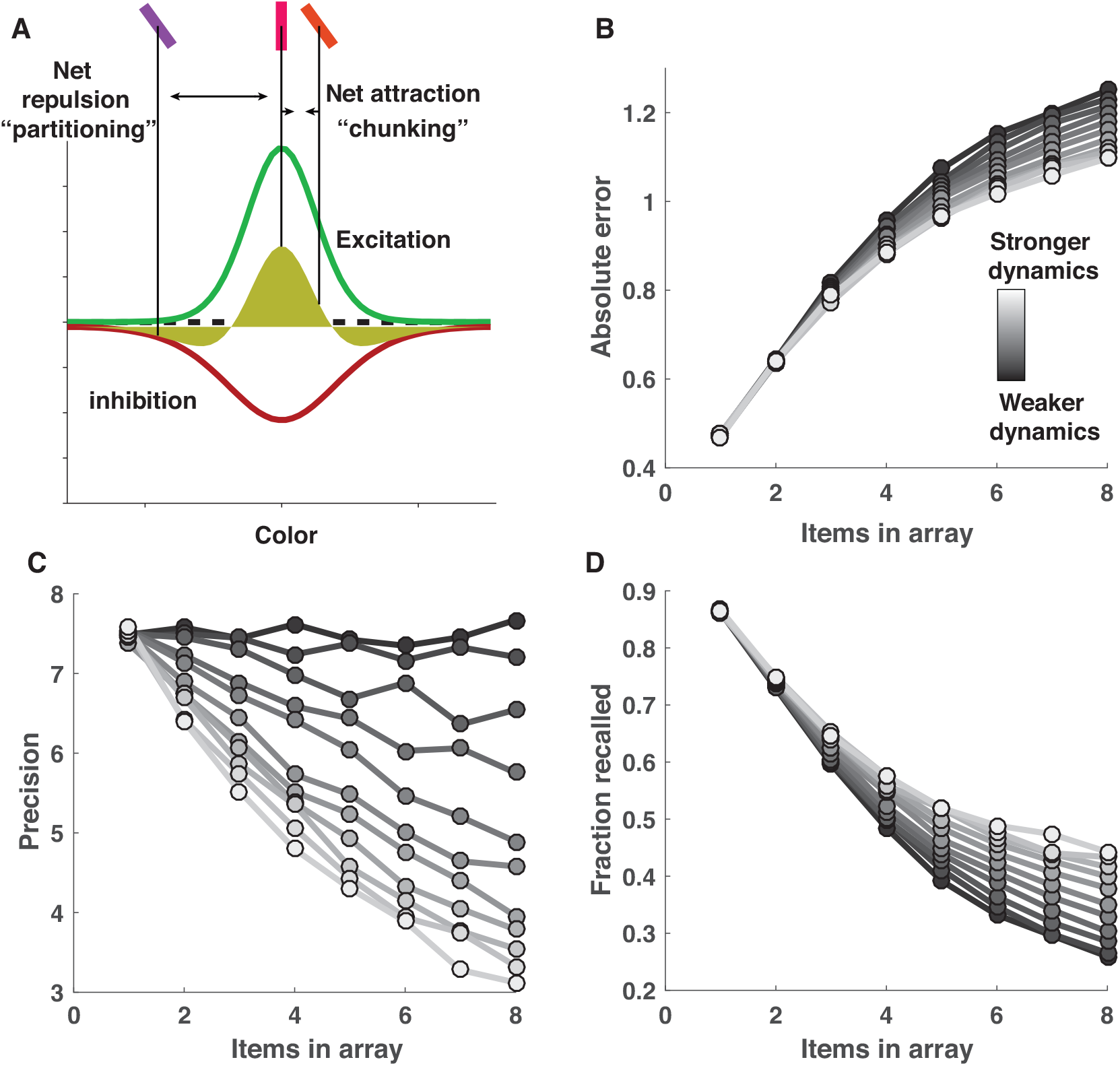
Center-surround dynamics facilitate attractive and repulsive inter-item forces that can improve recall at the cost of precision. **A)** Local recurrent excitation and broadly tuned lateral inhibition give rise to two counteracting forces: recurrent excitation facilitates attraction of neighboring representations through “bump collisions” (Wei et al., 2012), whereas broadly tuned lateral inhibition facilitates repulsion of distinct bumps of neural activity (Felsen et al., 2005; Kiyonaga & Egner, 2016). Together, these forces produce a difference of Gaussians tuning function (yellow shading) that facilitates attraction of closely neighboring representations but repulsion of more distant ones. Here we model these effects at the cognitive level by assuming that two imprecise internal representations of color are chunked, and jointly represented by their mean value, with a fixed probability defined by a narrowly tuned von Mises distribution (green curve; **B**&**C**) in order to mimic the effects of narrowly tuned excitation. After probabilistic chunking, each color representation exerts a repulsive influence over all other representations with a magnitude defined by a broadly tuned von Mises distribution (red curve) in order to mimic the effects of broadly tuned inhibition. The model stores a Poisson number of the representations, chunked or otherwise, for subsequent recall. **B**) The influence of center-surround dynamics over model performance can be manipulated by applying a gain to the amplitude of the excitation and inhibition functions such that larger values correspond to greater item interdependencies and lead to smaller errors on average (lighter colors correspond to higher gain). **C&D)** The performance improvement mediated by increasing center-surround dynamics relies on a tradeoff between recall probability and precision, through which increased attractive and repulsive forces reduce precision (lighter bars; **C**), but enhance recall probability (lighter bars; **D**).

We next examined how chunking, as implemented through this framework, would impact performance as a function of stimulus array size. To do so, we summarized the impact of center-surround connectivity in an algorithmic model of working memory performance that contained a small number of intuitive parameters. We implemented attractive and repulsive forces among stored memories in accordance with narrowly tuned excitation and broadly tuned inhibition functions (see Methods for details). On each trial, each color from the target array was: 1) perturbed by a mean zero random variable to simulate neural noise, 2) chunked with each other color in the array with probability proportional to the excitation tuning function, 3) repulsed by each other color in the array with magnitude proportional to the inhibition tuning function, and 4) probabilistically stored across the delay period according to a Poisson process. The proportionality constants allowed us to examine the performance of models ranging from those that were not affected by recurrent dynamics (zero proportionality constant) to those that were highly affected (large proportionality constant).

Models implementing greater recurrent dynamics achieved better performance through a recall/precision tradeoff. Performance was simulated on delayed report tasks in which target number (array size) ranged from one to eight. Performance of models employing recurrent dynamics was slightly worse for easier tasks but dramatically improved for more difficult ones, similar to the effects observed in the binary model above (figure 6B; lighter lines represent stronger recurrent dynamics). Here, though, performance differences were characterized by opposing effects of recurrent dynamics on precision and recall. Models employing recurrent dynamics showed improved recall, particularly in the hardest tasks, as attractive forces allowed for the storage of multiple target features in a single representation (figure 6D). However, these same recurrent dynamics came at the cost of reduced precision, as both attractive and repulsive forces reduced the fidelity of stored color representations (figure 6C). In standard models of resource limitations, precision decrements with increased array sizes have been attributed to the division of a limited resource. However, in the recurrent dynamics models, the decrement in precision is caused by the increase in inter-item interactions that occurs when additional items are added to the memory array. Thus, the inclusion of recurrent dynamics affects the nature of capacity limitations: minimizing the impact of center-surround forces leads to a specific decay in recall as a function of array size, as predicted by “slots” models, whereas maximizing the impact of center-surround forces leads to decays in precision across set size, which is a key feature of resource depletion accounts (Bays & Husain, 2008; Luck& Vogel, 2013; Ma et al., 2014; Zhang & Luck, 2008).

In summary, inter-item dependencies that emerge from center-surround dynamics are sufficient to mediate the performance bonuses of chunking, but do so at the cost of precision. Thus, if working memory is optimized through chunking in this way, it should lead to a higher probability of recall for colors of clustered stimulus arrays but more precise recall of colors for less clustered ones. In principle, such optimization could be guided in cognitive or real-world tasks by implicit or explicit feedback to favor successful chunking strategies and avoid unsuccessful ones.

### People are less precise when remembering clustered target arrays

Our center-surround implementation of chunking and partitioning predicts that chunking advantages should come at the cost of precision (figure 6B&C). To test this prediction, we examined the difference in error distributions for random vs. fixed spacing, pooled across all subjects (figure 7, left column). The error distributions from both conditions were consistent in shape with those previously reported in similar tasks (figure 7A&D) (van den Berg et al., 2014). However, the error distributions differed subtly between the two conditions: in the random-spacing condition, subjects made more moderately small errors, but did not have more perfect recalls (figure 7G). This pattern of difference was also evident in data simulated from the center-surround chunking model (figure 7, middle column) but not in data simulated from an independent encoding model fit to subject behavior (figure 7, right column). Thus, both the subjects and the center-surround chunking model reported more colors that were slightly offset from the target color in the random-spacing condition than in the fixed-spacing condition, consistent with a reduction in precision resulting from chunking.

**Figure 7:**
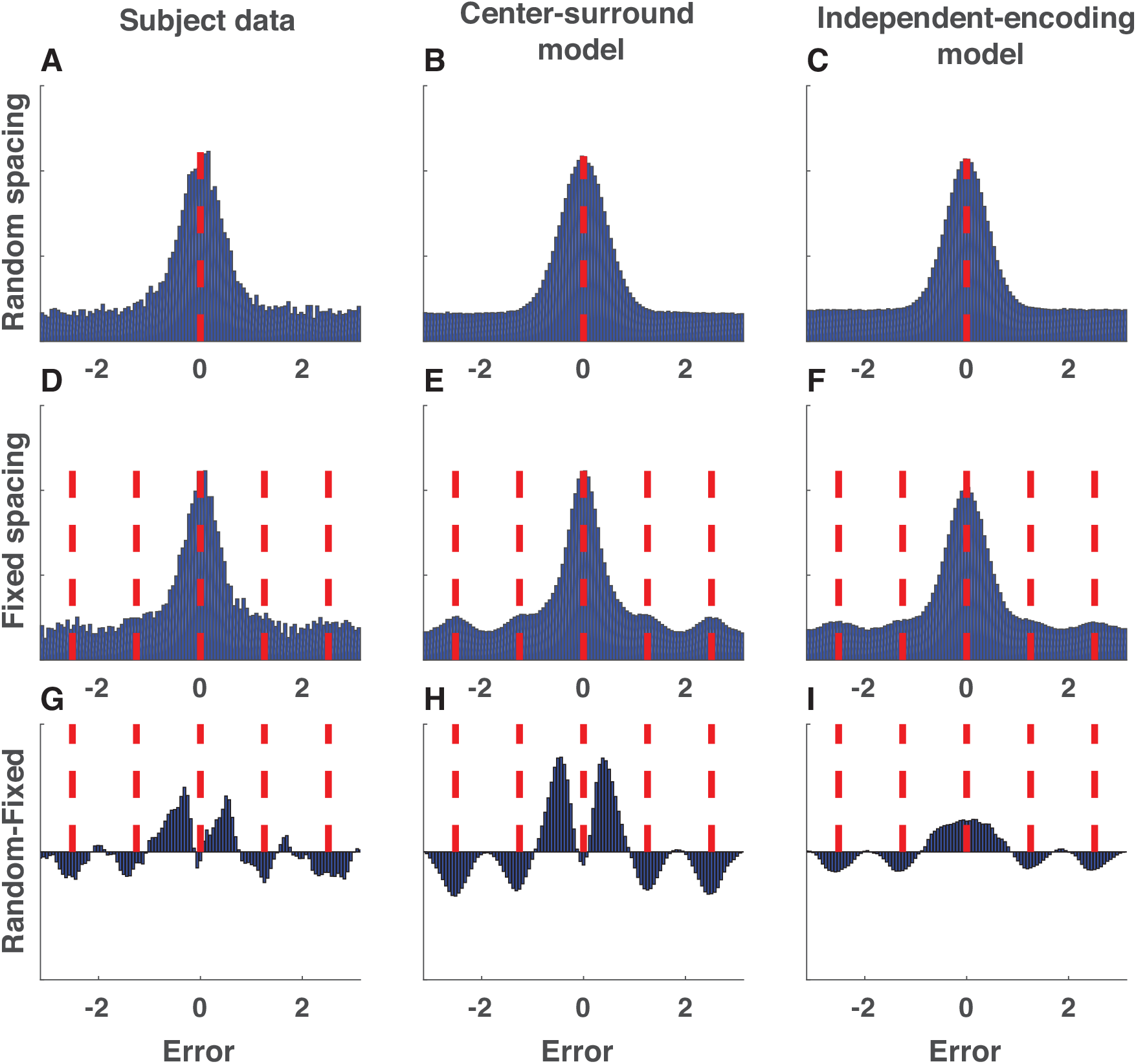
Error distributions reveal evidence for center-surround chunking. **A-C)** Signed color reproduction errors made in the random spacing condition by (**A**) subjects, (**B**) center-surround chunking models, and (**C**) independent encoding models. Data is collapsed across all simulated or actual sessions. **D-F)** Same as **A-C** but for the fixed spacing condition. Red dashed lines indicate probed and non-probed target locations. Note that the alignment of non-probed target locations emphasizes the prominence of non-probed target reports (binding errors), which would appear uniformly distributed in the random spacing condition. **G-I)** Difference in above error distributions for random minus fixed. To aid in visualization, bin count differences were smoothed with a Gaussian kernel (standard deviation = 1 bin). Subjects and the center-surround chunking model show increased moderately small, but non-zero, errors in the random spacing condition. Note that differences of reports between the random and fixed conditions near the non-probed targets are present in both models, as they simply reflect an artifact of the alignment of binding errors in the fixed spacing condition.

### Errors are modulated by nearest neighbors consistent with chunking via recurrence

To better understand the nature of these error distributions, and to what extent they are predicted by attraction and repulsion forces in center-surround dynamics, we sorted trials according to the non-probed target color that was most similar to the probed target color (nearest neighbor color; see Methods for details). This procedure revealed structure in individual subject color reports related to the nearest neighbor non-probed color (see figure S4). To determine whether such structure persisted systematically across subjects, we fit a descriptive mixture model to error distributions pooled across subjects in sliding windows of nearest neighbor distance. The model contained free parameters to examine 1) the precision of error distributions, 2) the bias of error distributions toward (or away from) the nearest neighbor non-probed target color, and 3) the relative proportion of trials that were recalled, forgotten, or mis-bound (in keeping with nomenclature from previous literature (Bays et al., 2009; Fallon, Zokaei, & Husain, 2016)).

The model fits revealed that subject precision and bias depended on neighboring colors in a manner consistent with chunking through recurrent dynamics. In particular, subject memory reports were biased towards the nearest neighbor color if it was sufficiently similar to the probed target color, but biased away from it if it was sufficiently dissimilar (figure 8A). This pattern of bias maps onto the idea of a narrowly tuned excitation function promoting attraction of nearby targets and a broadly tuned inhibition function promoting repulsion of more distant ones (see figure 8A). Precision also depended on nearest neighbor color distance: subject precision was maximal when the nearest neighbor color was most dissimilar to the probe color and minimal when it was moderately similar (figure 8D). In addition, fits revealed an increase in the proportion of correct recalls, and a corresponding decrease in the number of uniform guesses, when a nearby neighbor color existed in the stimulus array (figure S5). This pattern of results was strikingly consistent with those produced by a chunking model based on recurrent dynamics (figure 8, middle column) but not with those produced by the best-fitting mixture model (figure 8, right column).

**Figure 8:**
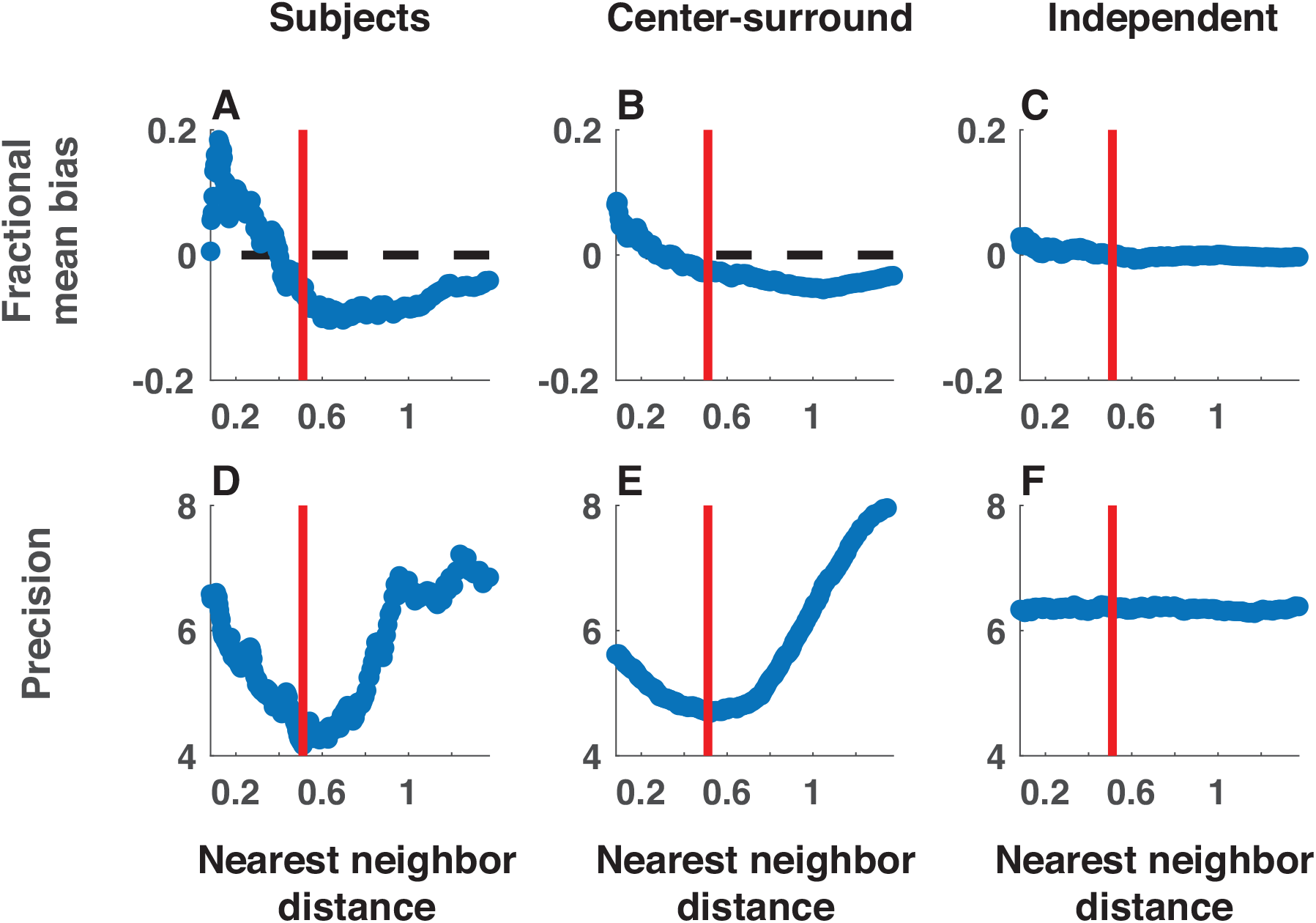
Neighboring stimulus features affect bias, precision, and recall probability as predicted by the center-surround chunking model. Subject (left) and simulated (center = center-surround, right = independent encoding) data were collapsed across all sessions and binned in sliding windows according to the absolute distance between the probed target color and the most similar non-probed target color (nearest neighbor distance; abscissa). Data in each bin were fit with *a* mixture model that included free parameters to estimate 1) the bias of memory reports towards the closest color in the target array expressed as a fraction of distance to that target (**A-C**), and 2) the precision of memory reports (**D-F**). The qualitative trends present in subject data are also present in data simulated from the center-surround chunking model but not in those simulated from the independent encoding model. Red bars reflect the nearest neighbor distance at which precision fits to subject data were minimal and also corresponds well with the crossover point of the bias fits.

### Quantitative model-fitting of center-surround model reveals empirical evidence of performance advantage for chunking

While the above mixture model fits revealed structure across subjects on average, here we provide a quantitative fit of the center-surround model directly to the trial-by-trial memory report data for each individual subject, allowing us to quantify chunking effects and examine the range of behavioral chunking strategies across individual participants. To make this fitting more tractable (i.e., to facilitate a closed form likelihood function), we simplified the center-surround model while retaining its core elements. These simplifications included removal of sensory noise and the simplification of the center (chunking) and surround (repulsion) forces (see Methods). We fit three different models to estimate the potentially separable contributions of center and surround functions using maximum likelihood. The “center” model estimated the partitioning criterion, which summarized the center function, as a free parameter, whereas the “surround” model fit the repulsion coefficient as a free parameter. The “center-surround” model fit both center and surround with free parameters. All models were also compared to a basic mixture model. Goodness of fit was evaluated for each model using AIC, which applies a fixed complexity penalty for each parameter and provided better model recovery for simulated data than BIC.

Comparison of the center-surround model to a basic mixture model revealed an explanatory advantage of the former, albeit with considerable heterogeneity across individual subjects. The center-surround model achieved the lowest mean relative AIC values of all models (mean[SEM] relative AIC: basic mixture = 4.9[0.9], center = 5.5[0.8], surround = 4.1[0.7], center-surround = 2.9[0.6]). Inclusion of both center and surround terms was favored by a likelihood ratio test (χ2(94) = 281, p <10e-5) and Bayesian model selection favored the center-surround model in terms of explaining the largest proportion of subject behavior (exceedance probability = 0.85). However, the best-fitting model was not consistent across subjects, with some subjects best fit by the simple mixture model and others best fit by the center-surround model (figure 9A). Parameter estimates from the best-fitting center-surround model were also indicative of heterogeneity. For a large number of subjects, the best-fitting partitioning criterions were near zero (indicating no chunking), but partitioning criterions fit to the remainder of subjects were broadly distributed (figure 9B). Best-fitting repulsion coefficients were more normally distributed across subjects, tending to take small positive values, indicating a tendency toward repulsion of partitioned representations (figure 9C).

**Figure 9:**
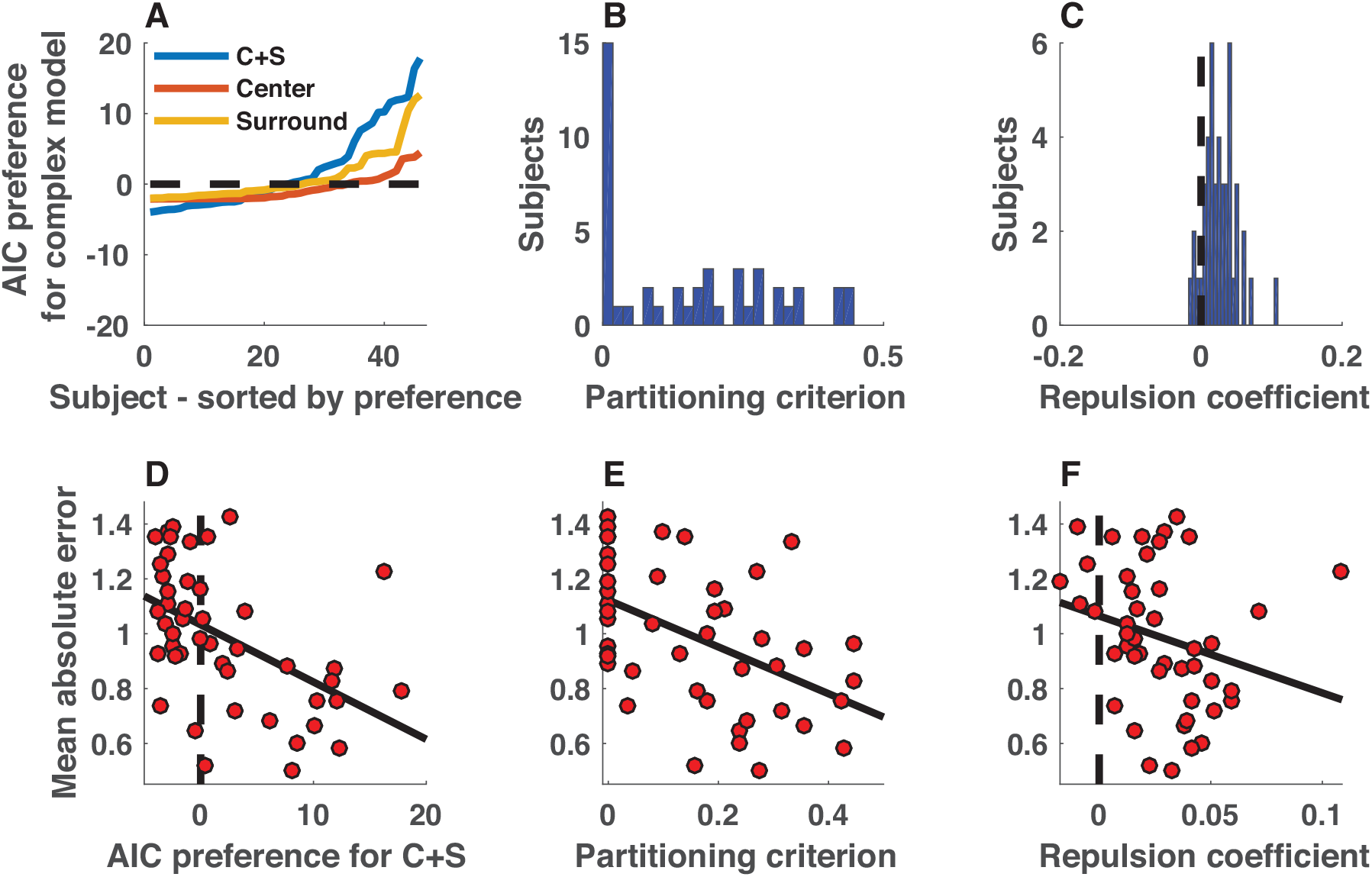
Heterogeneous chunking strategies across individual subjects provide empirical evidence for the performance advantages afforded by chunking. **A)** AIC difference between simple mixture model and more complex center (orange), surround (yellow), and center + surround (blue) models is plotted for each subject, sorted by model preference (positive values indicate that more complex model is preferred). Aggregate AIC values favored the C+S model, yet there was substantial variability across subjects in marginal improvement afforded by the C+S model over the simpler mixture model, with AIC values providing moderate evidence for the mixture in some subjects, but strong evidence for the C+S model in other subjects. **B)** Partitioning criterions best fit to subject data also reflected heterogeneity in strategies across subjects, with a number of subjects best fit with criterion values near zero, and another subset of subjects taking values across a wider range from 0.1-0.5. **C)** Best-fitting repulsion coefficients tended to take positive values across subjects, indicating that independently represented colors tended to exert repulsive forces on one another by the best-fitting model parameterization. **D-F)** Subjects displaying more evidence of center-surround chunking performed better on the working memory task. **D)** Mean absolute error was greatest for the subjects that displayed the least evidence of center-surround chunking, as assessed by the difference in AIC between C+S and basic mixture models (ρ = -0.59, p = 1.6e-5). **E&F)** Errors were also elevated for subjects that were best fit with criterions near zero (**E;** ρ = -0.54, p = 8.5e-5) or with small or negative repulsion coefficients (**F;** ρ = -0.39, p = 7.4e-3).

Heterogeneity in model fits also related to overall task performance in a manner suggestive of a performance advantage for chunking. Our modeling suggested that criterion-based chunking could be used to reduce overall errors in a visual working memory task, and the differences in model fits across our subjects offered us an opportunity to test this idea. Consistent with chunking facilitating in-task performance advantages, subjects fit with larger partitioning criterions and repulsion coefficients achieved better performance on the task (figure 9E&F; partitioning criterion: Spearman’s ρ = -0.54, p = 8.5e-5; repulsion coefficient: Spearman’s ρ = -0.39, p = 7.4e-3). Similar relationships were seen between model preference and overall performance, with the subjects that were best fit by the center-surround model also tending to have the lower absolute errors in the task (figure 9D; Spearman’s ρ = -0.59, p = 1.6e-5). In order to examine which parameter of our model best predicts subject performance, we constructed a GLM to examine the independent contributions of partitioning criterion and repulsion parameter estimates on subject performance (as measured by mean absolute error) and found that when accounting for both variables, only the partitioning criterion maintained explanatory power, with higher partitioning criterions corresponding to lower absolute error magnitudes (partitioning criterion β = -0.95, t= -3.2, p = 0.002; repulsion β = 0.8, t= 0.4, p = 0.7). Thus, individuals who chunked the most liberally also achieved the best task performance.

### Center-surround chunking effects generalize across tasks, contribute to set-size dependent changes in precision, and mediate individual differences in performance

Finally, to test whether our findings were robust to changes in task conditions and to examine how chunking effects vary with memory load, we fit a nested set of models to a meta-analysis dataset that included 101 subjects performing eight different experiments (James M Gold et al., 2010; van den Berg et al., 2014). The nested model set included models that varied in their assumptions about chunking, the distributional form of error reports, and the direct effects of set size on precision. The model set was built upon a base model that assumed that subjects would recall a Poisson number of feature representations in the report dimension and a Poisson number of features in the probe dimension on each trial, with failure to recall the probe dimension resulting in a binding error and failure to recall the report feature resulting in a uniform guess. Precision of memory reports was fixed across trials in this base model. The nested model set included additions to the base model that allowed it to account for 1) effects of center-surround chunking on the represented feature value and number of stored features (C in figure 10A), 2) effects of center-surround chunking on the precision of memory reports (N in figure 10A), 3) differences in error distribution kurtosis through t-distributed memory reports (T in figure 10A) and 4) changes in precision as a power-function of set size (P in figure 10A). Performance of the nested model set was compared to that of a variable precision model with Poisson item storage and binding errors, which was the best-fitting model in a recent factorial model comparison using most of the same data (VP in figure 10A, (van den Berg et al., 2014)).

**Figure 10:**
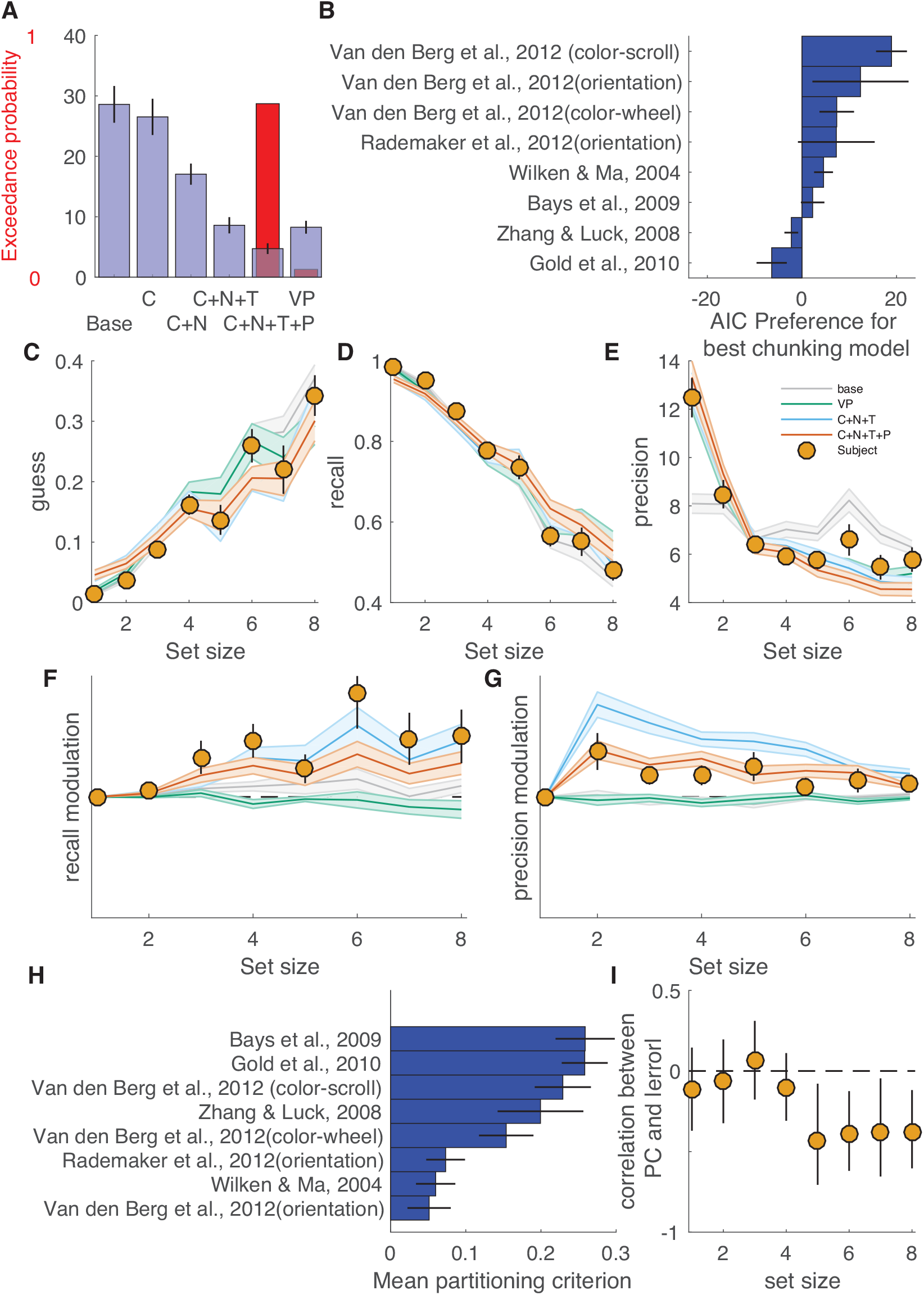
Center-surround chunking allows better fits of meta-analytic datasets and offers insight into trends and individual differences in how memory degrades with set size. **A)** Blue bars/lines represent mean/SEM relative AIC (AIC relative to that of the best model for each subject) and red bars reflect exceedance probability for a nested model set. Base refers to the base model, C includes center-surround chunking, N allows for chunking- and repulsion-induced report variability, T allows for t-distributed errors, and P allows precision to vary as a power-function of set size. Models are compared to the best-fitting model from a factorial model comparison that used this dataset (VP = variable precision, Poisson recall, with binding errors) (van den Berg et al., 2014). Bayesian model selection favored a model that incorporated t-distributed memory reports, power-law precision decrements and all modeled aspects of center-surround chunking (C+N+T+P). A model lacking power-law precision decrements (C+N+T) performed similarly in model comparison to the VP model. **B)** Horizontal bars reflect AIC preference for the winning (C+N+T+P) model over the best model that lacks chunking (VP) for each experiment and are arranged according to mean AIC preference (with experiments providing strongest support for the more center-surround model on top). **C-G**) Posterior predictive checks reveal nuanced discrepancies in the predictions across models. Actual and simulated data were sorted by subject and set size and fit with a flexible mixture model (see Methods) that estimated: guess rate (**C**), binding error rate (not shown), recall rate (**D**), report precision (**E**), modulation of recall by chunking (**F**) and modulation of precision by chunking (**G**). Points and lines reflect mean/SEM fits to subject data whereas lines/shading reflect mean/SEM fits to simulated data for each model (models denoted by color: base = gray, VP = green, C+N+T = blue, C+N+T+P = orange). All models captured guess and recall rates reasonably well (**C&D**), but only models that included either chunking (C+N+T), precision decrements with set size (VP) or both (C+N+T+P) could account for changes in precision of reports across set size (**E**). Only models that included chunking (C+N+T & C+N+T+P) could account for within set size modulation of recall (**F**). Within set size modulation of precision was overestimated by a chunking model with fixed assumptions about precision (C+N+T) and underestimated by models without chunking (base & VP) but well estimated by a model that included chunking and allowed precision to vary with set size (C+N+T+P). **H**) Bars indicate mean partitioning criterion for the (C+N+T) model across the experiments included in the meta-analysis (sorted from maximum). **I**) Correlation between mean absolute error magnitude (z-scored per experiment and set size) and the best-fitting partitioning criterion is plotted as a function of set size (abscissa). Points and lines reflect mean and bootstrapped 95% confidence intervals, respectively.

Model comparison favored the most complex model, which incorporated all aspects of chunking, flexible kurtosis of error distributions, and allowed precision to change as a function of set size (exceedance probability = 0.96, mean AIC relative to best AIC = 4.7; figure 10A). Performance of the model that included all aspects of chunking and kurtosis but not precision decrements was similar to that of the VP model (mean relative AIC 8.6 and 8.3 for C+N+T and VP models respectively; figure 10A). The advantage of the best performing chunking model was more prominent in some studies than others. The studies using the widest range of set sizes showed the largest chunking advantages (the Van den Berg et al. studies all included set sizes 1 through 8), whereas the study using the smallest range of set sizes (Gold et al, 2010; 3 or 4 items) showed a clear preference for the variable precision model over the best chunking model. It is worthy of note that some of the experiments included in the meta-analysis included additional manipulations or potential sources of variability that might have been captured by the variable precision model but could not possibly be accounted for in our chunking model, such as manipulation of the duration of stimulus presentation, the duration of delay, and the day of a multi-day experiment. Thus, despite the overall performance advantage of the best chunking model, it is still likely that some of the datasets include some residual variability that could be captured by additional variability in precision across conditions.

In order to better understand why the most complex chunking model offered a better fit to subject data, we performed a posterior predictive check by simulating data from each model using the maximum likelihood parameter values and then examining the simulated and actual data using an extended mixture model fit separately to data for each set size (figure 10C-G). While all models captured the general trends in recall and guessing (figure 10C-D), the base model was not capable of fitting the changes in precision across set size observed in subject data, whereas both the chunking models and variable precision model could capture these changes (figure 10E). Only the models that included chunking were capable of accounting for within-set size fluctuations in recall rate (figure 10F). Similarly, simulated data from both chunking models produced within-set size modulations of precision that were qualitatively similar to those observed in subjects, but the chunking model that lacked the ability to modulate precision according to set size produced much larger within-set size fluctuations in recall than were actually observed in the data. Thus, the best-fitting chunking model improved on the VP model by capturing the effects of center-surround chunking on recall and precision within set size. On the other hand, the chunking model that did not allow precision to change with set size seemed to capture set size precision effects by over-estimating the repulsive interactions between items, leading to a worse fit than the more complex model that allowed precision to change with set size.

Across experiments and subjects, there were systematic differences in chunking that related to overall performance. Mean partitioning criterion differed systematically across experiments in a manner unrelated to overall model preference (figure 10H). While the number of experiments included in this analysis is small, it should be noted that two of the experiments that included the least inclusive chunking behaviors involved storing an orientation rather than a color. The other experiment with a relatively small mean partitioning criterion (Wilken & Ma 2004) used a color sampling strategy that prevented fine grain estimation of the best-fitting partitioning criterion: similar but non-identical colors were never included in a single color array. Within experiments, subjects also differed in the partitioning criterion that best described their behavior in a manner that related to performance. Specifically, subjects that were best fit by the largest partitioning criterion values also tended to make the smallest absolute errors in the high, but not low, set size conditions (Figure 10I). Thus, individual differences in chunking may underlie individual differences in performance for high memory load conditions that have previously been thought to reflect differences in overall memory capacity.

## Discussion

Our work builds on two parallel lines of research. One has focused on how encoding and decoding of working memories are optimized under various statistical contingencies (Brady et al., 2009; Brady & Alvarez, 2011; Brady & Tenenbaum, 2013; Lew & Vul, 2015; Orhan & Jacobs, 2013; Sims et al., 2012), whereas the other has focused on understanding the nature of capacity limitations in visual working memory (Bays et al., 2009; Bays & Husain, 2008; van den Berg et al., 2012; 2014; Zhang & Luck, 2008; 2009; 2011). Here, we explore how people optimize encoding in the same tasks that have formed the basis of our understanding of capacity limitations. Our findings shed light on both the nature of memory capacity limitations and on the encoding strategies employed to minimize their impact.

With regard to encoding strategies, the binary encoding model showed that selective chunking allowed performance advantages for clustered stimulus arrays that grew as a function of set size, could be learned according to trial feedback, and limited asymptotic item storage to approximately four items (figure 3). Unlike previous models that have examined how non-independent item encoding and decoding schemes could affect memory performance (Brady et al., 2009; Brady & Alvarez, 2011; 2015; Lew & Vul, 2015; Orhan & Jacobs, 2013; Sims et al., 2012), our model shows how memory storage could be optimized without foreknowledge of, or even the existence of, statistical regularities in memoranda. Because of this, our model provides unique insight into how subjects might optimize behavior in standard working memory tasks, in which stimulus dependencies and foreknowledge thereof are intentionally minimized. As predicted by our model, human subjects display performance advantages when remembering clustered stimulus arrays (figure 4&S1) that are not explained by binding errors (Figure 4D&S2) and occur irrespective of whether the recalled item was itself in a stimulus cluster (Figure S3). Furthermore, trial-to-trial adjustments in performance of human subjects followed the same basic pattern through which chunking was learned in our model; namely, rewarded employment of chunking on one trial increased the probability of chunking on the next (Figure 4E&F). Thus, our model identifies and provides a normative explanation for a major source of performance variability across trials in visual working memory tasks: selective chunking of similar items into working memory and the optimization thereof.

Our findings are in line with previous work that highlights the use of chunking as a mnemonic strategy in a wide range of working memory tasks (Cowan, 2001). Chunking was first used to describe mnemonic strategies for storage of sequential information, for example, encoding the digits 2-0-0-5 as a single date (2005) rather than as its constituent digits (Chen & Cowan, 2005; Cowan, 2001; G. A. Miller, 1956). In the visual domain, visual features are in some sense chunked into objects (Luria & Vogel, 2014). Recent work has suggested that people can chunk arbitrary visual information when that information is inherently clustered and visible for an extended duration (Lew & Vul, 2015). Here, we extend on this work to show that a simple form of chunking, joint encoding of similar feature values, is rapidly implemented by human visual working memory systems to improve performance in tasks that have heretofore been thought to lack exploitable statistical structure.

The basic computations necessary to achieve performance advantages through chunking could be endowed to a recurrent neural network by implementing center-surround dynamics (figure 5). These dynamics arbitrate a tradeoff between recall and precision (figure 6) that was supported by empirical evidence of higher precision representations for unclustered stimulus arrays (figure 7). This sort of tradeoff between memory precision and item capacity is similar to that observed in binding pool models of working memory, where the level of connectivity in the binding pool controls a tradeoff between precision and quantity of representations (Swan & Wyble, 2014; Swan, Collins, & Wyble, 2016). Here we show that this tradeoff can be exploited to improve performance, and that human subjects seem to do so. In particular, subjects demonstrated the performance benefits, response biases, and costs in precision that were predicted by center-surround chunking and were quantitatively best described by it (figures 5&7–9). While the quantitative advantage for the center-surround model was small (figure 9), this advantage was likely limited in part by technical constraints that required simplification of the center-surround model for fitting purposes, in particular by the removal of noise in the initial color representation. Nonetheless, these findings are generally consistent with previous work that has highlighted the effects of center-surround processing on perception and memory (Almeida et al., 2015; Johnson, Spencer, Luck, & Schöner, 2009; Störmer & Alvarez, 2014; Xing & Heeger, 2001). Furthermore, the specific inter-item dependencies predicted by our model and observed in our empirical data were consistent with those emerging from recurrent neural networks that rely on tuned inhibition (Almeida et al., 2015), but not with those predicted by hierarchical models of memory decoding, as the latter do not produce repulsion of dissimilar features (Brady & Alvarez, 2015; Orhan & Jacobs, 2013).

Our center-surround model serves not only to describe nuanced features of behavior, but also to link our findings to potential biological mechanisms. We show that a small change to a prominent neural network model of working memory maintenance, namely the incorporation of tuned inhibition, provides the model with the capability to chunk similar features into a joint representation but partition dissimilar ones through repulsion (figure 5; (Ben-Yishai et al., 1995; Kohonen, 1982; Murray et al., 2014; Somers et al., 1995; X. J. Wang, 1999; Wei et al., 2012)). Our descriptive model based on this mechanistic account is supported by the frequent observation of sustained activity during the delay period of memory tasks in both parietal and prefrontal cortices (Funahashi, Bruce, & Goldman-Rakic, 1989; Fuster & Alexander, 1971; Gottlieb, 2004) (but see also (Lara & Wallis, 2014)). Here we have considered the network to store features on a single dimension (color); however, it is clear that at some level, conjunctive coding across features (e.g. color and orientation) is necessary to bind information to the dimension used to probe memories (Matthey, Bays, & Dayan, 2015). In our task, it is unknown whether any sustained representations reflect information about the report feature (color in our task), probe feature (orientation in our task), or some conjunction of the two. Recent work has hinted that in some cases, sustained representations in prefrontal cortex may only encode the probe dimension, which could point back to relevant sensory representations at time of recall (Ester, Sprague, & Serences, 2015; Kriete, Noelle, Cohen, & O’Reilly, 2013; Lara & Wallis, 2014; 2015). Previous computational instantiations of this process have relied on the basal ganglia to learn appropriate prefrontal representations that can be jointly cued by multiple disparate perceptual features, based on reward feedback (A. G. E. Collins & Frank, 2013; Frank & Badre, 2012; Kriete et al., 2013). Analogously, feedback effects observed in our data could be driven by the basal ganglia learning to selectively engage prefrontal units that are prone to representation of multiple probe feature values. This interpretation could expand on a large body of work that implicates the basal ganglia in gating working memory processes by stipulating a novel and testable role for the basal ganglia in optimizing joint feature encoding (Chatham, Frank, & Badre, 2014; A. G. E. Collins & Frank, 2013; Hazy, Frank, & O’Reilly, 2006; O’Reilly & Frank, 2006; Voytek & Knight, 2010).

An important question stemming from our work is to what extent chunking can be adjusted to optimize working memory accuracy under different conditions. Our modeling shows that a simple learning rule is capable of rapidly adjusting the amount of chunking to optimize performance given the current memory demands, leading to greater chunking for higher memory loads. Individual differences in chunking were selectively related to performance in the highest memory load conditions (figure 9E & 10I); however, neither our experiment, nor the meta-analytic dataset explicitly manipulated chunking incentives over a time-course long enough to measure learning effects. Nonetheless, even in the absence of explicit manipulation, feedback-dependent modulation of chunking behaviors in our experimental data was indicative of online optimization of the chunking process (figure 4D-F), such as the process that allowed learning of the partitioning criterion in the binary encoding model (figure 4G). Yet these trial-to-trial adjustments occur despite only minimal performance improvements across task blocks (figure S2). There are several possible explanations for this discrepancy, including 1) that a priori processing strategies are well-calibrated to our task, 2) that optimization in our task occurs on a different timescale than our measurements, and 3) that the presence of uniformly spaced arrays hindered learning overall. Distinguishing between these possibilities will require a better understanding of what exactly is being adjusted in response to feedback. For example, reward feedback could promote the prioritization of storing chunked arrays over non-clustered ones, or it could modulate center-surround inhibition dynamics (e.g., via fine tuning of feature selective attention and/or altering local excitation-inhibition balance (Störmer & Alvarez, 2014; Wei et al., 2012)). In any case, our work, along with other recent research showing an adaptive tradeoff of precision and recall (Fougnie, Cormiea, Kanabar, & Alvarez, 2016), strongly motivate future work to better understand the scope, time course, and mechanism for this type of optimization process.

### Implications for capacity limitations

Working memory limitations have been theorized to result from either a discrete limitation on available “slots” or a continuous limitation by a divisible “resource”. The distinction between these theories is most evident when additional targets are added to a memory array. A discrete limitation predicts that after all slots are filled, additional targets will be forgotten and will be reported as random guesses (Luck & Vogel, 2013). In contrast, a resource limitation predicts that additional targets will cause each target to be encoded with lower precision (Ma et al., 2014). While individual studies have provided support for each theory (Bays et al., 2009; Bays & Husain, 2008; Cowan & Rouder, 2009; Chris Donkin et al., 2013a; Christopher Donkin et al., 2013b; Pratte, Park, Rademaker, & Tong, 2017; Rouder et al., 2008; van den Berg et al., 2012; Zhang & Luck, 2008; 2009; 2011), a recent meta-analysis provides simultaneous support for the core predictions of both: increasing memory load leads to both increased guessing and decreased precision (van den Berg et al., 2014).

Our results suggest that a joint capacity limitation over recall and precision may result in part from a rational chunking procedure implemented through center-surround dynamics to effectively trade precision for recall (figure 5&6). This procedure allows subjects to achieve performance improvements for clustered stimulus arrays at the cost of precision (figures 6). It is also capable of explaining decrements in precision with set size, as larger sets of items lead to increased repulsive forces experienced by each individual item (figure 10E). In addition to accounting for known influences on precision, our model also predicted that measured precision should vary across trials, an established feature of human behavioral data (Fougnie et al., 2012; van den Berg et al., 2012), and correctly predicted that this variability in precision should depend on the features of non-probed targets (figure 8D&E; 10G). Nonetheless, the best-fitting center-surround chunking model employed leptokurtic memory reports in order to capture additional variability in precision that was not accounted for by the chunking and repulsion processes alone, suggesting that other factors must also contribute to variability in precision (figure 10A). Furthermore, the best-fitting model also allowed memory report precision to vary as a power function of set size, as this appropriately balanced the magnitude of across set size (figure 10E) and within set size (figure 10G) precision effects. Thus, center-surround chunking, as we implemented it, can quantitatively account for most, but not all, of the changes in precision across trials and set-sizes.

Our findings also call the interpretation of precision measurements into question. The center-surround model predicts that internal representations apply attractive and repulsive forces to one another, systematically biasing memory reports. When averaged across trials with differing stimulus configurations, such interactions are interpreted as variability in memory reports, as the net forces on a probed target vary randomly from one stimulus configuration to the next. Yet, since much of this variability is simply an artifact of averaging across disparate conditions, our work raises an important question: how much of the variability in memory reports across trials and individuals is truly reflective of imprecision, rather than bias? While the notion that imprecision can emerge from systematic inter-item dependencies is somewhat at odds with the basic resource limitation model, it is consistent with the recent proposal of a specific form of resource limitation in which the constrained resource is the representational space itself (M. A. Cohen, Rhee, & Alvarez, 2016; Franconeri, Alvarez, & Cavanagh, 2013; Oberauer & Lin, 2017).

Within such a framework, it is interesting to reconsider the meaning of individual differences in memory storage recall and precision. Previous work has shown that individual differences in the number of items successfully retained in visual working memory tasks, but not differences in precision, are related to fluid intelligence and psychiatric conditions such as schizophrenia, among other factors (Fukuda, Vogel, Mayr, & Awh, 2010; James M Gold et al., 2010). These results have been interpreted in terms of differences in a discrete capacity for memory storage, or in filtering irrelevant information (Vogel, McCollough, & Machizawa, 2005), but our results suggest that some of these individual differences may be driven instead by differences in chunking behavior or the optimization thereof. To this effect, we showed both in our own dataset and in the meta-analytic dataset that individual differences in task performance, particularly at high set sizes, were systematically related to differences in chunking policy, with subjects that chunked most liberally achieving the best performance for higher set sizes (figures 9E&10I). Given that the best chunking policies in our binary encoding model for a set size of five were quite liberal (figure 3A,C,D,F) we suspect that a number of subjects could have improved performance dramatically, were they to have chunked more liberally. In fact, a subset of subjects was best fit by models that fully partitioned color information (figure 9A), achieved lower overall performance (figure 9E), and likely attenuated the aggregate performance advantages for clustered stimulus arrays (compare figure 3L to figure 4A). It is not clear to what extent these individual differences in chunking policy result from differences in the ability to learn an appropriate criterion, or from hard-wired differences that might predispose individuals toward either chunking or partitioning neural representations. Our neural network model suggests that performance-based measures of capacity may be sensitive to individual differences in lateral connectivity profiles that favor a spectrum from independent to merged feature storage policies, and to the ability to override such policies through learned top-down modulation of lateral connectivity (Freeman, Driver, Sagi, & Zhaoping, 2003; Freeman, Sagi, & Driver, 2001; Lowe et al., 2016).

In summary, our results show that humans readily exploit chunking strategies to improve performance on visual working memory tasks. The implementation of chunking is consistent with a form of center-surround dynamics that combines similar representations and facilitates mutual repulsion of disparate ones. This implementation leads to a fundamental tradeoff between the number of items stored and the precision with which they are stored, providing a natural bridge between slots and resource accounts of working memory capacity limitations. People optimize this tradeoff from trial-to-trial according to stimulus statistics and evaluative feedback in a manner that differs across individuals and is predictive of working memory task performance. These results provide a normative joint account of how and why discrete and continuous factors contribute to working memory capacity limits across individuals and task conditions.

## Methods

### Delayed report task

54 human subjects completed five blocks (100 trials each) of a delayed report color reproduction task (figure 2). Each trial of the task consisted of four primary stages: stimulus presentation, delay, probe, and feedback. During stimulus presentation, subjects were shown five oriented bars (length = 2 degrees visual angle) arranged in a circle (radius = 4 degrees visual angle) centered on a fixation point. Bar positions were equally spaced around the circle and jittered uniformly from trial to trial. Bar orientations were uniformly spaced, jittered from trial to trial, and independent of position or color. Bar colors were chosen from a fixed set of colors corresponding to a circle in CIELAB color space (L= 80, radius in a*, b* = 60) and referred to by angular position for convenience. In the “random spacing” condition, all five colors were sampled independently of one another from the color space, allowing for the possibility of two similar colors in the same stimulus array. In the “fixed spacing” condition, colors were uniformly spaced along the CIELAB color wheel and randomly assigned to bar locations. Stimuli were presented for 200 ms, after which the screen was blanked.

The subsequent delay period lasted 900 ms, during which subjects were forced to remember the colors and orientations of the preceding stimulus array. During the subsequent probe stage, subjects were shown a gray oriented bar in the center of the screen for one second, before being asked to report the color that had been associated with that orientation in the preceding stimulus array. Color reports were made by adjusting the color of the oriented bar using a mouse. The initial position of the mouse on the color wheel was randomly initialized on each trial. On a subset (1/3) of trials, subjects were asked to make a post-decision wager about the accuracy of their report by choosing to bet either 0 or 2 points. Binary feedback was provided on each trial based on whether subject reporting accuracy fell within a certain error tolerance window (*π* /3 radians – low precision condition [26 subjects] or *π* /8 radians – high precision condition [28 subjects]). A priori target sample size for each group was set to twenty-four based on other studies in the field (without explicit power calculations). Additional subjects were recruited beyond this to account for potentially unusable data (e.g. subjects guessing on all trials). Four subjects in the high precision condition and three subjects in the low precision condition were removed from analyses because of error distributions that were not statistically distinguishable from uniform guessing (error variance > 0.91), leading to sample sizes of 23 and 24 for low and high precision conditions respectively. All subjects were paid bonuses according to total accumulated points. All human subject procedures were approved by the Brown University Institutional Review Board and conducted in agreement with the Declaration of Helsinki.

### Binary encoding model

To explore the potential advantage of chunking in delayed report tasks, we developed a flexible and computationally tractable model for capacity-limited storage. This model stores color and orientation information symbolically in a set of binary “words” concatenated to form a “sentence”. During the stimulus presentation phase, target colors and orientations are “encoded” as an alternating sequence of binary words reflecting the position on a circular feature space (figure 3). The number of binary digits (bits) in a word controls the precision with which the feature is stored. For example, a single digit can encode which half of the feature space contains the color of a bar, whereas three bits can narrow the stimulus color down to one eighth of the color space (figure 3, top). Each binary word is followed by a “stop” symbol denoting the type of information in the preceding word (e.g. color or orientation). A capacity limitation is implemented in the model as a limit on the number of bits that can be stored in memory. Specifically, we applied a fixed limit of 15 bits for storage of color information. Similar results were achieved by applying a limit to the total bits, i.e. including orientation information, but here we allow for perfect orientation storage in order to isolate the effects of capacity limitations on the recall dimension (color).

Bits were allocated in two different ways: in one set of simulations, bits were assumed to be continuously divisible (analogous to resource models) and in the other set of simulations, bits were not divisible beyond binary units (analogous to slots-and-averaging models). For resource model simulations, performance was computed analytically according the following error function:

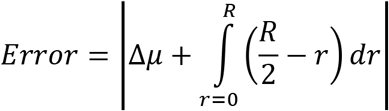

where Δμ is the difference in the “chunk” mean and the true color and R is the continuous range over possible stimulus values specified by the encoding model, which depends on the number of bits allocated to each item according to the following function:

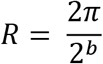

where b is the number of bits allocated to each target, which in turn depends on the total number of items and the exact pattern of chunking and partitioning across the stimulus values. Chunking and partitioning were controlled in three ways: 1) in the fully partitioning model, all colors were represented separately, 2) in the optimal partitioning model, all possible partitioning patterns were considered for each stimulus array and only the performance of the best partitioning pattern was reported, and 3) in the criterion-based partitioning model, adjacent colors were partitioned (represented distinctly) if they were separated by a distance that exceeded the partitioning criterion. Colors that were not separated by partitions were combined into a single “chunk” and represented by their mean value. This procedure could sometimes lead two colors that were separated by distances greater than the partitioning criterion to be included in the same chunk, if they were both sufficiently close to an intermediate color. The performance of the criterion-based partitioning model was computed across a range of possible partitioning criterions and the performance of the best-performing criterion across all trials for a given set size was reported (see figure 4A).

In the second set of simulations where bits were considered to be indivisible, analogous to the slots + averaging framework, model performance was assessed through exhaustive simulations. In this framework, bits were as evenly distributed among represented colors as was possible for a given stimulus array, as this strategy for allocation of bits achieved the best performance. During the probe phase, the model is presented with a single orientation and recalls the color word that immediately precedes that orientation in the stored binary sentence. A report is then sampled from a uniform distribution across the range of colors consistent with that stored binary color word. For example, if the color word contains one, two, or three bits, it is sampled from uniform distribution over one half, quarter, or eighth of the color space.

Chunking was parametrically implemented in the binary encoding model by adding a “partitioning criterion” that specifies the minimum distance between two colors in color space that is necessary for independent storage. Colors separated by distances greater than the partitioning criterion are partitioned, and all colors that are not separated by a partition are combined into a single color representation. The distance computation is completed during the “encoding” phase, before colors are converted to binary words. Distances are corrupted with a small amount of independent noise consistent with variability in the visual representation or the chunking processes (normally distributed with standard deviation equal to 0.4 times the partitioning criterion). This noise gave rise to variability in the chunking process, such that a given set of stimuli might be partitioned differently on different trials. After chunking, bits are allocated evenly across all represented colors, as described above.

Model performance was simulated for the delayed estimation task across eight different array sizes (1-8) with two different color generation conditions (fixed- and random-spacing) for nine different partitioning criterions ranging from zero to π. For each condition and model, mean absolute error was computed across 5000 simulated trials. The best chunking model (see figure 4L) was defined as the model with the lowest mean absolute error, whereas the fully partitioned model was the model with partitioning criterion equal to zero (such that every color was stored independently). For each condition, chunking bonus was computed as the difference in absolute error between the non-chunking and best-chunking models.

For the trial-to-trial optimization of the partitioning criterion (figure 4g), we adjusted the partitioning criterion on each trial according to the following rule:

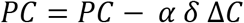

where PC is the partitioning criterion, *α* is a learning rate, δ is a reward prediction error (previous trial feedback minus long term average feedback), and *ΔC* is the number of “chunks” into which the previous stimulus array was divided minus the long term average of that quantity. Thus, if by chance the model did more chunking on a given trial, the *ΔC* would take a negative value, and positive feedback would drive a positive *δ* and a corresponding increase in the partitioning criterion, leading to an increase in chunking on subsequent trials. Negative feedback for the same trial would lead to a negative δ and corresponding decrease in the partitioning criterion, leading to a decrease in chunking on subsequent trials.

### Computing array clustering

In order to assess the potential benefits of chunking on each trial we computed a clustering statistic, within-cluster variance (WCV), for each stimulus array. WCV was computed by dividing the array colors into two clusters that minimized the mean variance within the clusters. WCV was defined as the average circular variance over colors within these clusters.

### Logistic regression models

Binary accuracy and betting data were concatenated across all subjects and interrogated with a mixed-effects logistic regression model that included terms to account for fixed effects of 1) -log(WCV), a proxy for stimulus array chunkability, 2) the color distance between the probed target and each other color in the array, ordered from smallest to largest, 3) feedback on previous and subsequent trials, 4) spatial distance between the location of the probed target and the location of the previously probed target, and 5) task block. In addition, the model included dummy variables to account for random intercepts specific to individual subjects. The same analysis was applied to data simulated from the best-fitting mixture model, which considered all reports to come from a weighted mixture of recall, uniform guess, and binding error distributions (Bays et al., 2009).

### Mixture model

We extended the standard mixture model of memory reports (Bays et al., 2009; Zhang & Luck, 2008) to allow for modulation of recall probability, precision, and bias according to WCV, nearest neighbor distance, and feedback. The standard mixture model assumes reports are generated from a mixture of “correct recall”, “guessing”, and “binding error” processes. These three mixture components were specified using two free parameters: one dictating the probability with which an item would be successfully stored *(correct recall* + *binding error)* and one specifying the probability with which a stored item would be correctly reported 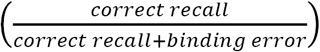. We allowed the parameter dictating successful storage to be modified as a logistic function of 1) log(WCV), 2) previous feedback, 3) previous log(WCV), **4**) previous feedback*log(WCV) and 5) previous feedback*log(WCV)*previous log(WCV). All potential modulators of successful storage were mean-centered (before and after interaction) and constrained by priors favoring values near zero [~normal(0, 0.5)]. Since our successful storage parameter is the probability the subject will not elicit a uniform guess, it affects both correct recall and binding error mixture components. However, since reports were far more likely to correspond to correct recalls (median mixture proportion = 0.50 across subjects) than binding errors (median binding error proportion = 0.17 across subjects), modulator coefficients had larger effects on recall than binding errors, and we refer to them in the results as modulating recall for simplicity. We also considered an alternative model in which modulators affected the recall term directly and found similar results, although this alternative model provided a worse overall fit of the data. Group mean parameter estimates were tested against the null hypothesis (estimate = 0) with a classical one sample t-test; however, in cases where moderate p values were observed (p <0.1 & p > 0.01) Scaled-Information Bayes factors were computed to quantify the model evidence in favor of the alternative hypothesis relative to that of the null hypothesis (rscale=0.707) {Rouder:2009ij}.

### Neural network simulations

Neural network simulations were conducted using a basic recurrent neural network that has been described previously (Wei et al., 2012). The model consists of 2048 pyramidal (excitatory) neurons and 512 inhibitory interneurons. Pyramidal neurons had the following cellular properties: C_m_ = 0.5 nF, g_leak_= 0.025 μS, V_leak_ = -70 mV, V_thresh_ =-50 mV, V_res__=-60 mV, τ = 1 ms. Interneurons had the following cellular properties: C_m_ = 0.2 nF, g_leak_= 0.02 μS, V_leak_ = -70 mV, V_thresh_ =-50 mV, V_res__=-60 mV, τ = 1 ms. The model included AMPA, NMDA, and GABA receptors with properties described previously (Furman & Wang, 2008). Pyramidal-to-pyramidal connection weights followed a narrowly tuned Gaussian profile across stimulus space (σ =5, J^+^ =5.6). Pyramidal-to-interneuron and interneuron-to-pyramidal connectivity profiles were identical, and in one set of simulations fully connected with uniform weights (figure 5C). In the second set of simulations, the cross-population connectivity was defined by a mixture of uniform weights and broadly tuned Gaussian weights (σ =20, mixture proportion = 0.1). Input was delivered to both networks for 200 ms through activation of an AMPA current with g_max_ = 0.57 using a spatial profile that was centered on 5 “target colors” with a Gaussian profile (σ =4). Stimulus delivery was followed by a delay period during which no input was provided to the network and activity was sustained completely through recurrent connectivity.

### Center-surround chunking model

To determine the effects that center-surround dynamics would have on visual working memory task performance, we extended the standard descriptive model of delayed memory reports to incorporate features of center-surround dynamics. In particular, on each trial, internal representations of each color were generated from a von Mises distribution with fixed concentration (7 for simulations). Pairwise distances (in color space) were computed for each pair of internal representations. Chunking probability was computed as a scaled von Mises function of this distance (μ = 0, κ = 12 for simulation), corresponding to the narrow excitatory “center” over which local representations are likely to attract one another (figure 5A-C). Representations were merged in accordance with these chunking probabilities by replacing the color associated with each merged representation with the mean of the merged colors. After probabilistic chunking, distances were recomputed between representations, and each representation applied a repulsive force on neighboring representations as defined by a scaled von Mises function of the re-computed distance (μ = 0, κ = 2 for simulation), corresponding to the broadly tuned “surround” over which representations repulse one another (figure 5A-C). Applying these forces leads each representation to be reset according to the following equation:

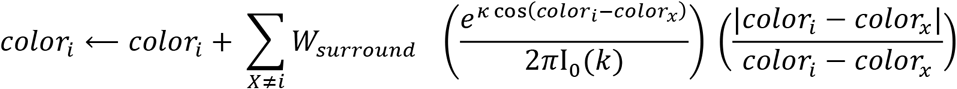

where *W_surround_* is a weight that controls the overall magnitude of surround effects, the second term in the sum is the probability density function for a von Mises distribution, and the final term serves to ensure that targets exert repulsive forces on neighboring targets. For the simulations in figure 5, the weight parameters for both center and surround were set to equal values ranging from 0 to 0.7. For comparisons to subject data, *W_surround_* was set to 0.6 and *W_center_* was set to 1.2.

Probabilistic recall was implemented in the model according to a Poisson memory process (Sims et al., 2012; van den Berg et al., 2014). On each trial, the model accurately recalled some number of representations drawn from a Poisson distribution (λ = 2 for simulations). Similar results were achieved using an inhibition based forgetting process inspired by Wei and colleagues (Wei et al., 2012); however, here we use a more standard Poisson process for simplicity. In the case that a representation that was not successfully recalled was probed, the model reported either a uniformly distributed guess (p = 0.65) or the color of an alternative representation (binding error, p = 0.35).

### Quantitative model fitting

In order to estimate model-likelihood directly, the center-surround chunking model was modified to allow for a closed-form likelihood function. To this end, we stipulated that internal representations would perfectly reflect the true stimulus colors before being subjected to chunking and repulsion processes instead of assuming that internal representations were subject to variability resulting from perceptual processing (as described above). In order to improve gradient descent, we implemented chunking using a gamma distribution over partitioning criterions in which the mean of the distribution was fit as a free parameter and the variance was fixed to 0.01. The repulsion process was simplified to a linear function of inter-item similarity, with a slope that was fit as a free parameter and could take either positive values to capture attraction or negative values to capture repulsion. Three versions of the simplified chunking model were fit to delayed report data: 1) a center-only model in which the partitioning criterion mean was fit as a free parameter and the repulsion coefficient was fixed to zero, 2) a surround-only model in which the partitioning criterion mean was fixed to zero and the repulsion coefficient was fit as a free parameter and 3) a center-surround model in which both terms were fit as free parameters. In addition, all models included the following free parameters: 1) Poisson lambda to describe the number of items that would be stored on a given trial, 2) binding error fraction to describe the frequency that reports would be generated from a non-probed representation, and 3) precision of the report distribution. All models were compared to a basic mixture model (Bays et al., 2009) using to penalize for complexity, as AIC allowed for better model recovery from simulated data than did BIC. AIC values are reported for each model relative to the lowest AIC model achieved by any model for a given subject (relative AIC). Bayesian model selection was performed using -1/2 AIC as a proxy for model evidence with the SPM toolbox (Stephan, Penny, Daunizeau, Moran, & Friston, 2009).

### Meta-analysis

In order to test the robustness of our findings and determine how the behavioral hallmarks of chunking scale with the size of the stimulus array, we applied a modified version of our mixture model to a meta-analysis dataset. The meta-analysis dataset included eight studies and a total of 101 subjects (Bays et al., 2009; James M Gold et al., 2010; Rademaker, Tredway, & Tong, 2012; van den Berg et al., 2012; Wilken & Ma, 2004; Zhang & Luck, 2008). Seven of the datasets, available online at http://www.cns.nyu.edu/malab/resources.html, were originally compiled by van den Berg et al. and have previously been described in detail (van den Berg et al., 2014). Three of the studies compiled by Van den Berg et al. were excluded from our analyses due to retraction of the original studies, although the inclusion of these studies did not qualitatively change our results. The eighth dataset (28 subjects) comprised the control subjects in a psychiatric comparative study of visual working memory (James M Gold et al., 2010). Each study differed in experimental details but involved a delayed report working memory task with at least two different array sizes.

### Quantitative model fitting to meta-analytic data

We constructed a nested set of models to better understand whether chunking could improve explanations of behavior in previous studies visual working memory studies. Each model was extended beyond a “base” model in which the partitioning criterion was fixed to zero. The base model included one change from the models that were fit to the data from our experiment in order to account for the possibility that binding errors depend on set size (which was variable in the meta-analytic data but fixed in our own study). Specifically, we replaced the fixed-probability of binding errors with a Poisson distribution that described the number of probe dimension features that would be recalled on a given trial (*s*), with lambda of this distribution fit as a free parameter. For each trial, this distribution was used to compute a probability that the relevant probe feature would not be stored in memory:

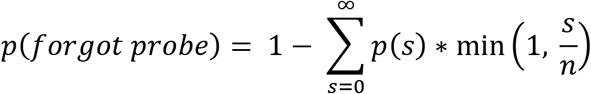

where *n* is the number of targets presented on a given trial and p(*s*) is the probability of recalling *s* probe dimension features on a Poisson distribution. The minimum term accounts for the case where the number of available items is smaller than the number of probe dimension features that could have been stored on a given trial.

The probability of making a binding error, given that the recall feature was remembered, was then computed as:

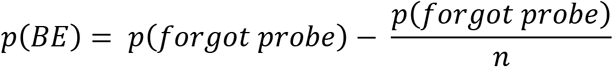

to correct for the possibility that, in the case that the model did not correctly store the probe dimension feature, it could choose the correct report dimension feature by chance. This change allowed the model to capture tendencies for binding errors to increase with set size, as have been reported previously (Bays et al., 2009).

The first extension to the base model allowed the partitioning criterion and repulsion criterion to be fit as free parameters, rather than set to zero as they were in the base model. This extension allowed the model to capture the biases and recall benefits that are predicted by our more general center-surround chunking model (e.g. figures 5–8); however, it would not capture variability in reports that would be expected to occur through the amplification of sensory noise by the chunking and repulsion processes, as the sensory noise was removed in order to allow for a closed form likelihood function.

In order to account for the basic effects of chunking and repulsion on report variability that would be expected based on our center-surround chunking model, but maintain the tractability of our likelihood function, we added two additional parameters to the model to allow the variance in memory reports to scale linearly with 1) the variance of feature values stored within a single chunk [chunking noise], and 2) the total repulsive forces experienced by the recalled feature [repulsion noise].

We also considered an extension that employed a more flexible report distribution that included an additional free parameter to model differences in kurtosis. In this extension, memory reports were generated from a t-distribution centered on the value of the internal representation and truncated at that value plus or minus π radians. The t-distribution included a base scale parameter fit to each subject, which accounted for overall variability in memory reports and was incremented by the additional chunking and repulsion variability as described above. In addition, the t-distribution included a degrees of freedom parameter that was fit to each individual subject, which allowed the model to capture report distributions ranging from leptokurtic (low degrees of freedom) to mesokurtic (high degrees of freedom).

Finally, we considered an extension to the model that included the possibility that precision depends on set size. Specifically, we stipulated that precision, or inverse variance, of memory reports would obey a power-law relationship with set size:

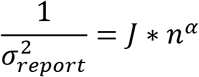

where *J* is the response precision expected when set size is equal to 1, and α is the power delineating the dependency of precision on set size, with negative values of α corresponding to precision values that decay with set size.

The nested set of models were tested against one another, and also compared to a variable precision model that includes Poisson item limits and binding errors that scale linearly with set size, which was the best performing model in a previous meta-analysis of delayed report working memory behavior (van den Berg et al., 2014). Model comparison using AIC and Bayesian model selection was done as described above. Posterior predictive checks were conducted by fitting actual and model-generated meta-analytic data with a descriptive model of memory report distributions separately for each subject and set size. The descriptive model estimated the rate of three response types (guess, binding errors, and correct recall) and the precision of memory reports as has been described previously (Bays et al., 2009). However, the model also included two additional terms to capture fluctuations in recall and precision within a given set size that would be predicted by chunking. Specifically, the model allowed the probability of recall to vary as a logistic function of trial-to-trial recall probabilities extracted from the center-surround chunking model, and allowed the variance of the report distribution to vary as a linear function of the trial-to-trial prediction for response variance extracted from the center-surround chunking model. Trial-to-trial model predictions were extracted from the center-surround chunking model that included chunking and repulsion noise as well as t-distributed errors, as this model provided a combination of a good fit to most subject data and relatively well-behaved parameter estimates.

### Nearest neighbor analysis

For each trial, the nearest neighbor color was identified as the color of the non-probed target that was most similar to that of the probed target. Target colors and subject reports were transformed for each trial such that the probed target color corresponded to zero and the nearest neighbor color ranged from zero to π. Trials were then sorted according to absolute nearest neighbor distance (see figure S2) and binned in sliding windows of 50 trials according to nearest neighbor distance. Binned data were combined across all subjects and fit with a mixture model that assumed data were generated from a mixture of 1) a von Mises distributed memory report (free parameters: mean, precision, and mixture weight), 2) uniformly distributed guesses (free parameters: mixture weight), and 3) binding errors that were von Mises distributed and centered on non-probed targets (no free parameters required, as mixture weight forms simplex with the other mixture components). Maximum posterior probability parameter estimates for the mixture model fits to subject and model simulated data are reported in figure 9 (prior distributions for all modulator terms were centered on zero with σ = 0.5 for recall modulators, σ = 2 for precision modulators, and σ = 0.05 for bias modulators).

## Author Note

We thank Ronald van den Berg, Wei Ji Ma and Jim Gold for sharing data from previously published delayed report studies. We thank Xiao-Jing Wang for providing code for neural network simulations. We thank Karen Schloss for help with accurate color rendering. We thank Anish Aitharaju, Anthony Jang, Ji Sun Kim, Michelle Kulowski, and Ezra Nelson for aiding in data collection. We thank Nicholas Franklin and Nathan Vierling-Claassen for helpful discussion. This work was funded by NIMH grant F32 MH102009 and NIA grant K99AG054732 (MRN), as well as NIMH grant R01 MH080066-01 and NSF grant #1460604 (MJF). A previous version of this work was presented at the New England Research on Decision Making (NERD) conference and was published as a preprint on biorxiv: http://biorxiv.org/content/early/2017/01/06/098939.

## Supplementary Information

**Figure S1:**
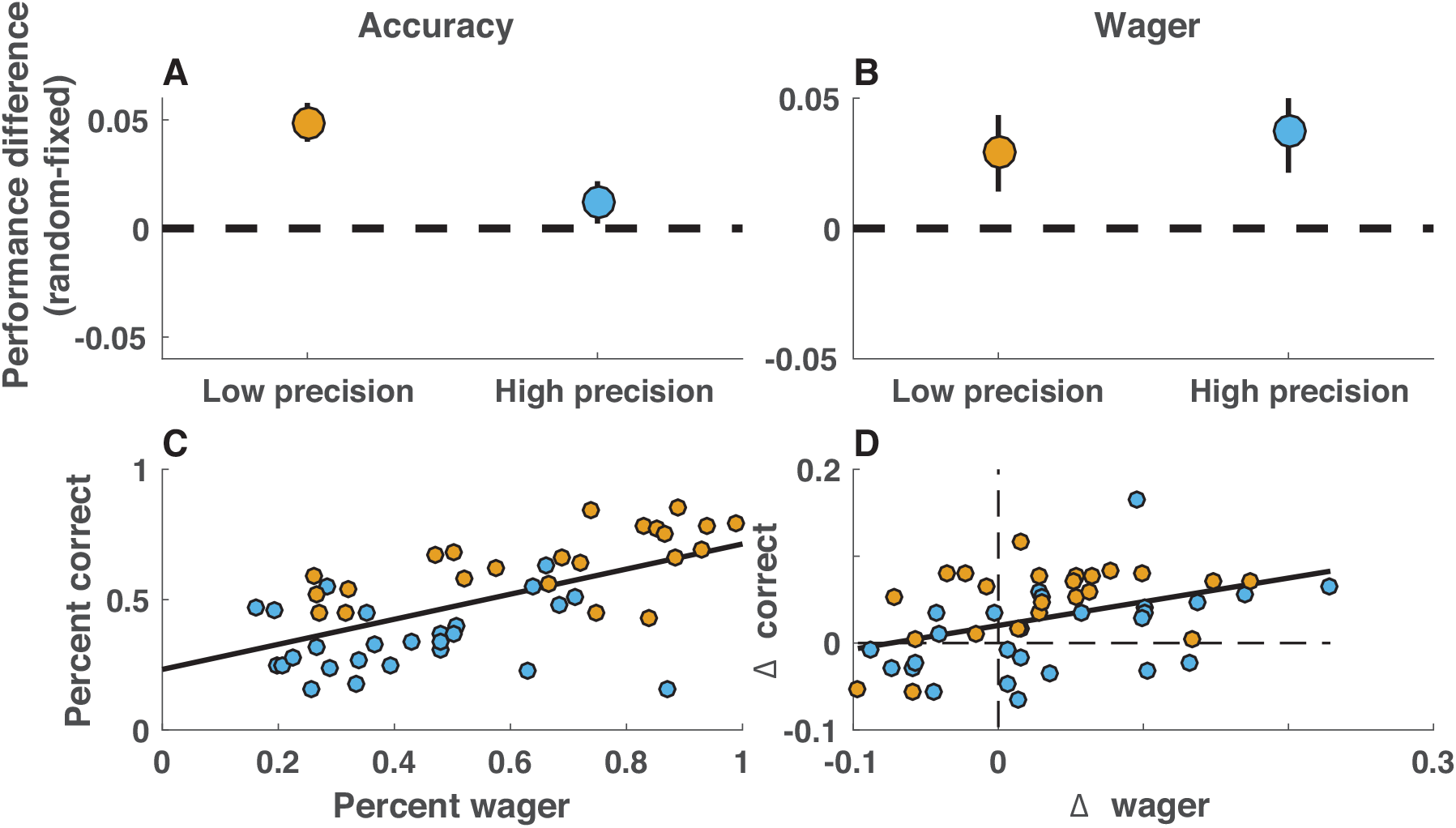
Uniformly spaced stimulus configurations degrade task performance and confidence in human subjects. Subject performance was assessed in terms of accuracy (percent of trials eliciting positive feedback) and confidence (percent of trials eliciting high post-decision wagers) separately according to precision condition (23 subjects were required to achieve an error of less than π/3 to elicit positive feedback [low precision], whereas 24 subjects were required to achieve an error of less than π/8 to elicit positive feedback [high precision]). **A)** Subjects in the low precision condition were more accurate for random spacing, as opposed to fixed spacing, stimulus configurations (orange; t=5.6, p < 10e-4), whereas subjects in the high precision condition attained similar overall performance in both configurations (blue; t=1.5, p = 0.15). Points/lines indicate group mean/SEM. **B)** Subjects in both conditions indicated higher confidence for random-spacing, as opposed to fixed-spacing, stimulus configurations (t = [2.3, 2.0] and p = [0.03, 0.06] for high and low precision conditions, respectively). **C)** Subjects that were most accurate, as assessed online according to a fixed error threshold, also tended to make higher post-decision wagers. Orange and blue points indicate subjects in low and high precision conditions, respectively. **D)** Furthermore, the improvement in accuracy from fixed- to random-spaced arrays was greater for subjects that showed the largest increase in confidence across the same conditions.

**Figure S2:**
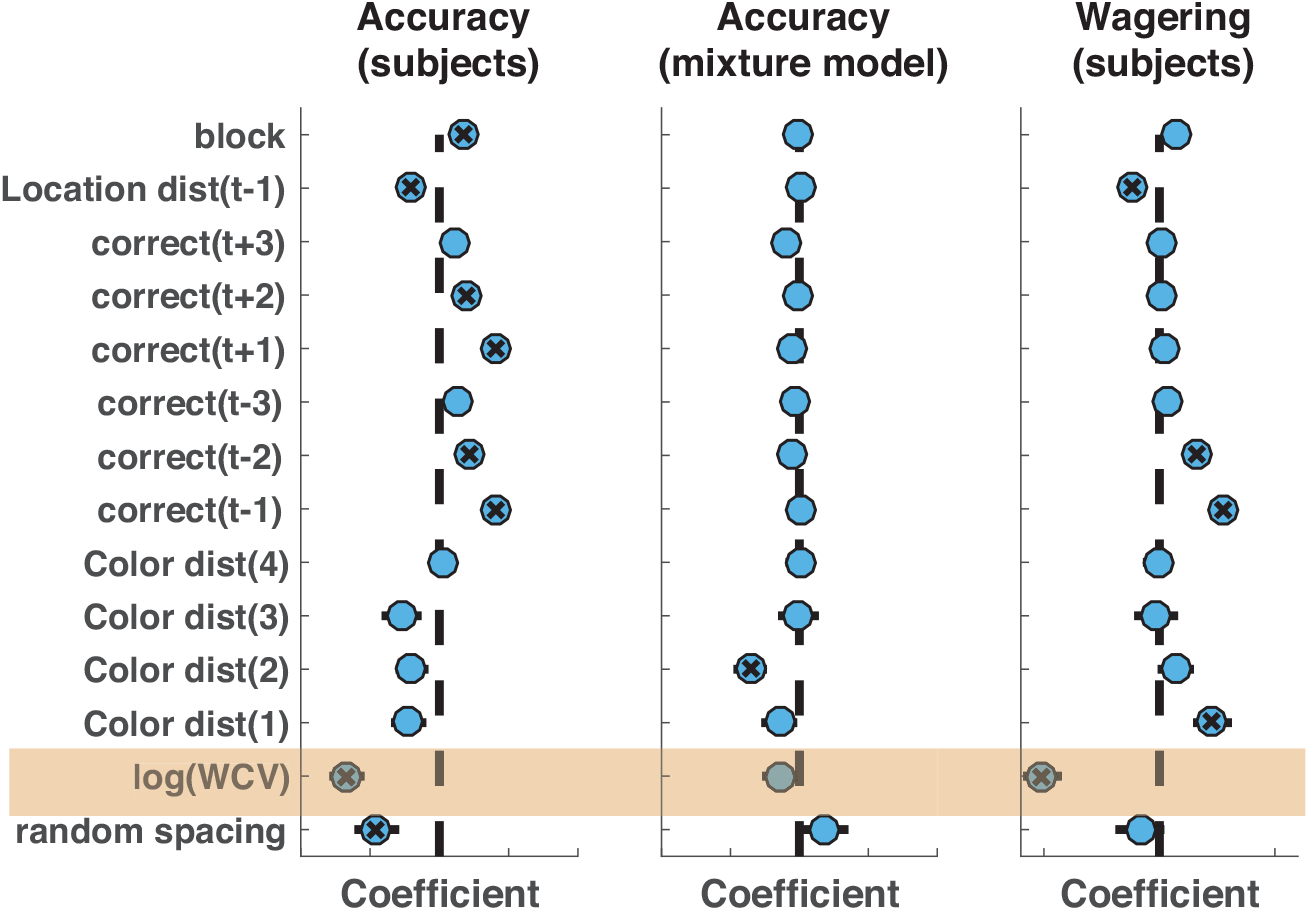
Stimulus clustering and recent feedback impact accuracy and confidence after controlling for other confounded factors in a GLM. Effects of color clustering on performance (left) and confidence (right) persist after accounting for potential confounding factors and feedback-dependent performance adjustments. Coefficients from a mixed-effects logistic regression model of binary accuracy and wager are plotted on the abscissa. Circles/lines reflect mean/SEM, and X marks indicate coefficients significantly different from zero (p < 0.05). Coefficients for log(WCV), a proxy for stimulus clustering, are highlighted in orange. Coefficients for log(WCV) are significantly lower than zero, indicating better performance / higher wagering on trials where stimulus colors were more clustered. This effect is not present in accuracy data simulated from a mixture model that includes binding errors (center).

**Figure S3:**
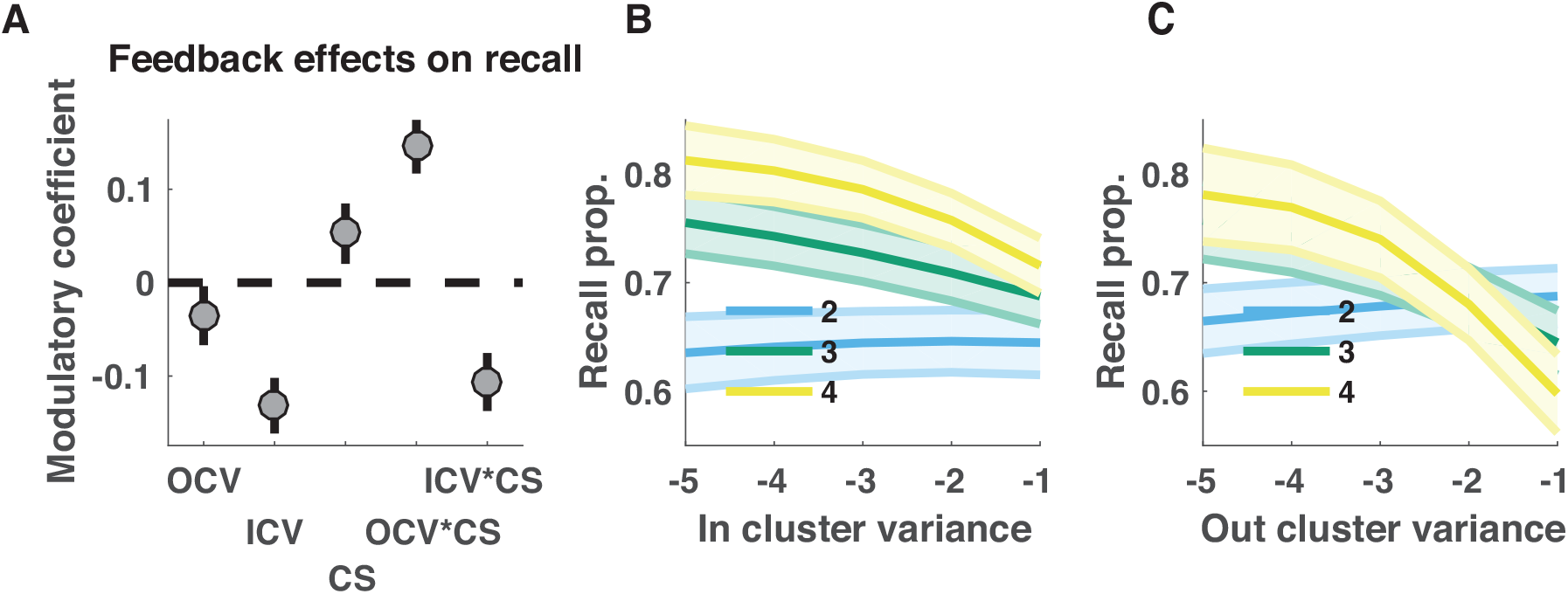
Recall is affected by clustering of both probed and non-probed stimuli. To further examine the source of the recall benefits, subject data were fit with a mixture model that considered reports to come from a mixture of processes including 1) a uniform “guess” distribution, 2) a “memory+binding” distribution centered on the color of the probed target, and 3) a “binding error” distribution including peaks at each non-probed target. Additional terms were included in the model to allow the recall probability to vary as a logistic function of various descriptive aspects of stimulus clustering that all factor into the within-cluster variance measurements reported in figure 5. To do so, the color array from each trial was divided into two (minimal variance) clusters to compute 1) the variance of the cluster that did not contain the probed item [OCV], 2) the variance of the cluster that did contain the probed item [ICV], and the number of colors in the cluster that contained the probed target [CS]. **A)** Mean/SEM coefficients across subjects indicated that these three factors, along with their interactions, were systematically related to trial-to-trial fluctuations in subject recall rates. **B&C)** The predicted recall rates from model fits are plotted as a function of ICV (B) and OCV (C) color coded according to the number of items in the relevant cluster (the cluster containing the probed item for B, and the cluster that did not contain the probed item for C). Recall bonuses are evident for low values of both ICV and OCV, although these benefits scale with the number of colors contained in the relevant cluster.

**Figure S4:**
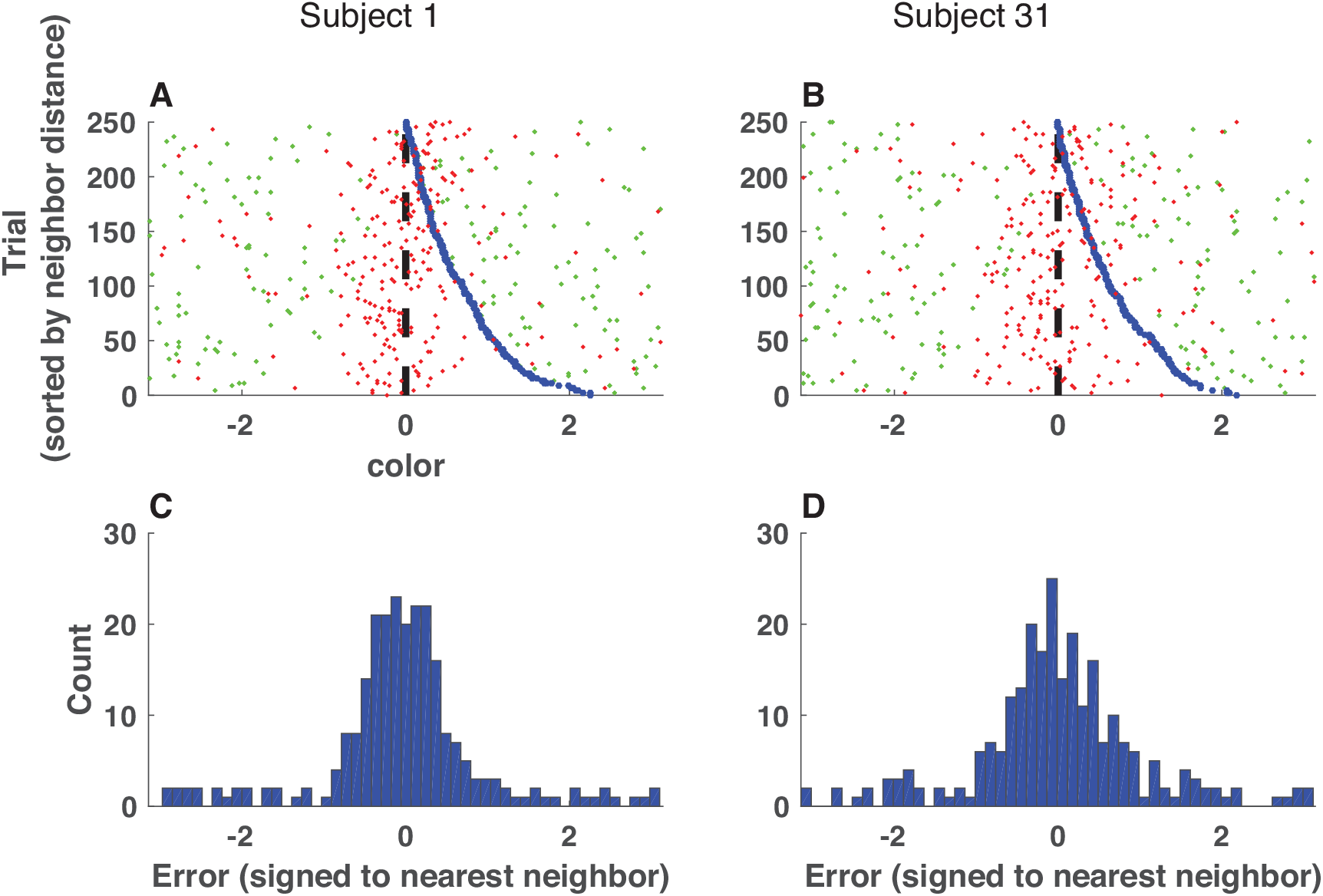
Sorting trials according to the nearest neighbor non-probed target color reveals structure in memory reports. **A&B:** Signed error of memory reports (red points) for all trials completed by two sample subjects (left = subject 1, right = subject 31). Trial errors are sorted by the distance from the probed target to the most similar color in the target array (nearest neighbor distance, NND) and transformed according to the direction of the nearest neighbor target (blue points). Green points reflect the positions of other colors in the target array, relative to the probed color and transformed as described above. Note the asymmetry in error distributions appears to change as a function of the nearest neighbor distance. **C&D:** Error histograms for the same two example subjects, transformed as described above. Note that in some cases apparent structure in the sorted errors (A) is no longer visible after collapsing across nearest neighbor distances (C).

**Figure S5:**
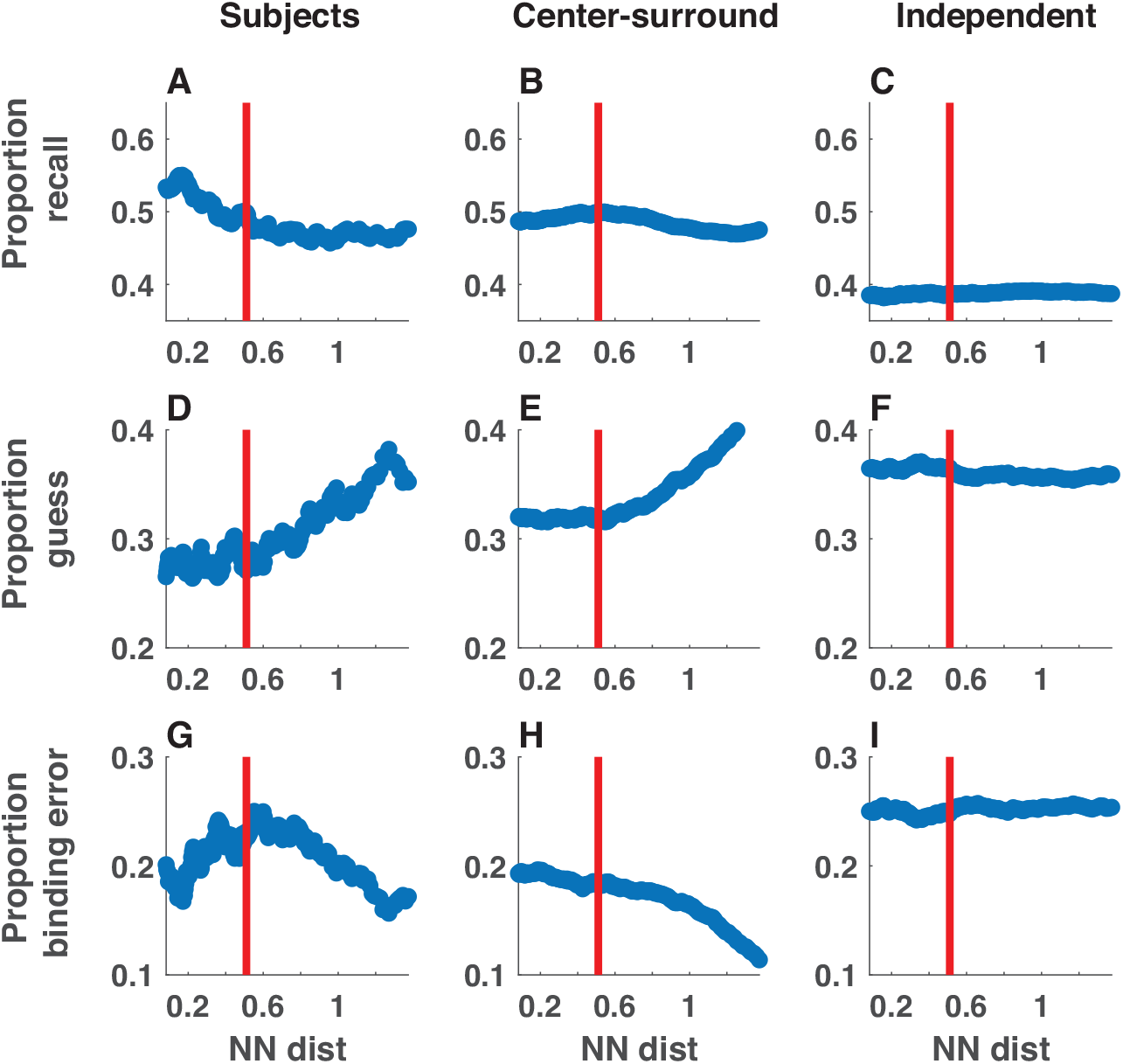
Neighboring stimulus features affect fits of mixture model. Subject (left) and simulated (center = center-surround, right = independent encoding) data were collapsed across all sessions and binned in sliding windows according to the absolute distance between the probed target color and the most similar non-probed target color (NN dist; abscissa). Data in each bin were fit with a mixture model that included free parameters to estimate the proportion of reports generated from 1) the von Mises “memory distribution” (**A-C**), 2) the uniform “guess distribution” (**D-F**), or 3)the mixture of von Mises “binding error distribution” (**G-I**). Parameter estimates for precision and bias terms are reported in the main text (figure 8).

